# DNA-loop extruding SMC complexes can traverse one another *in vivo*

**DOI:** 10.1101/2020.10.26.356329

**Authors:** Hugo B. Brandão, Zhongqing Ren, Xheni Karaboja, Leonid A. Mirny, Xindan Wang

**Affiliations:** Graduate Program in Biophysics, Harvard University, Cambridge, MA 02138, USA; Department of Biology, Indiana University, Bloomington, IN 47405, USA; Institute for Medical Engineering and Science, Massachusetts Institute of Technology, Cambridge, MA 02139, USA

**Keywords:** SMC complexes, loop extrusion, Z-loops, condensin

## Abstract

The spatial organization of chromosomes by structural maintenance of chromosomes (SMC) complexes is vital to organisms from bacteria to humans ^1,2^. SMC complexes were recently found to be motors that extrude DNA loops ^3–11^. It remains unclear, however, what happens when multiple SMC complexes encounter one another *in vivo* on the same DNA, how encounters are resolved, or how interactions help organize an active genome ^12^. Here, we set up a “crash-course track” system to study what happens when SMC complexes encounter one another. Using the *parS*/ParB system, which loads SMC complexes in a targeted manner^13–17^, we engineered the *Bacillus subtilis* chromosome to have multiple SMC loading sites. Chromosome conformation capture (Hi-C) analyses of over 20 engineered strains show an amazing variety of never-before-seen chromosome folding patterns. Polymer simulations indicate these patterns require SMC complexes to traverse past each other *in vivo*, contrary to the common assumption that SMC complexes mutually block each other’s extrusion activity ^18^. Our quantitative model of bypassing predicted that increasing the numbers of SMCs on the chromosome could overwhelm the bypassing mechanism, create SMC traffic jams, and lead to major chromosome reorganization. We validated these predictions experimentally. We posit that SMC complexes traversing one another is part of a larger phenomenon of bypassing large steric barriers which enables these loop extruders to spatially organize a functional and busy genome.

## Main text

Chromosomes from all kingdoms of life are actively maintained and spatially organized to ensure cell viability. SMC complexes play a key role in spatially organizing chromosomes and function in many processes like chromatin compaction, sister-chromatid cohesion, DNA break repair, and regulation of the interphase genome ^1,2^. While their importance has been recognized for over 25 years, evidence for a molecular mechanism of how SMC complexes function has only recently emerged. Recent single molecule experiments and chromosome conformation capture (Hi-C) studies have shown that the condensin and cohesin SMC complexes can translocate on DNA and extrude DNA loops at rates of ~1 kb/s ^3–11^. The process of DNA loop extrusion by SMC complexes is emerging as a universal mechanism by which these proteins organize the 3D genome in eukaryotes and prokaryotes ^2^. It remains unclear, however, what happens in a living cell when multiple SMC complexes encounter one another ^19^. Understanding the outcome of such encounters is fundamental to elucidating how chromosomes are spatially organized by the process of DNA loop extrusion.

Encounters between SMC complexes are expected to occur frequently in a cell. In eukaryotes, the cohesin and condensin SMC complexes are loaded at multiple chromosomal loci, at estimated densities between ~1/200 kb - 1/40 kb and extrude loops of 100’s of kilobases (reviewed in ^18^). In many bacteria, condensin SMC complexes are loaded by the protein ParB primarily at centromeric sequences called *parS* sites ^13–17^. These sites often exist in multiple copies close to one another ^20^. In bacteria without the ParB/*parS* system, such as *E. coli,* the condensin-like MukBEF complex loads non-specifically, but creates long DNA loops ^21,22^. Therefore, in both eukaryotes and bacteria, SMC complexes will frequently encounter others when extruding DNA loops. Most efforts towards understanding the chromosome organizing capacity of SMC loop extruders have assumed that translocating complexes are impenetrable to each other ^23^ (also reviewed in ^2,19^). A recent single-molecule study using *Saccharomyces cerevisiae* condensins challenged this assumption and demonstrated that condensins can traverse past each other *in vitro* ^12^. How SMC complexes interact *in vivo, (i.e.* traversing, blocking, or unloading each other, etc.), and the implications of these interactions for chromosome folding remain unknown. Here we show that *B. subtilis* condensins can traverse past each other *in vivo* in a quantitatively predictable manner, resulting in an unexpected diversity of chromosome folding structures.

We set up a condensin complex “crash-course track” system to probe the effects of encounters between loop extruding factors (Fig. 1A). We engineered *B. subtilis* strains to contain one, two or three *parS* sites, and we varied the relative separations and positions of the condensin loading sites (Fig. 1B). This allowed us to better resolve the effects of encounters between condensins than in the wild-type system, which has nine *parS* sites in proximity to one another ^24–26^. Moreover, to remove potential confounding effects of interactions between the replication machinery and condensins, and to eliminate potential interactions between sister chromatids, we synchronized cells in G1 phase by expressing the protein SirA. SirA inhibits replication initiation while allowing ongoing rounds of replication to complete, leaving cells with single chromosomes ^27^. We then investigated chromosome interaction patterns using Hi-C and protein distributions by Chromatin Immunoprecipitation (ChIP-seq) assays ^3,25,28^.

**Figure 1:**
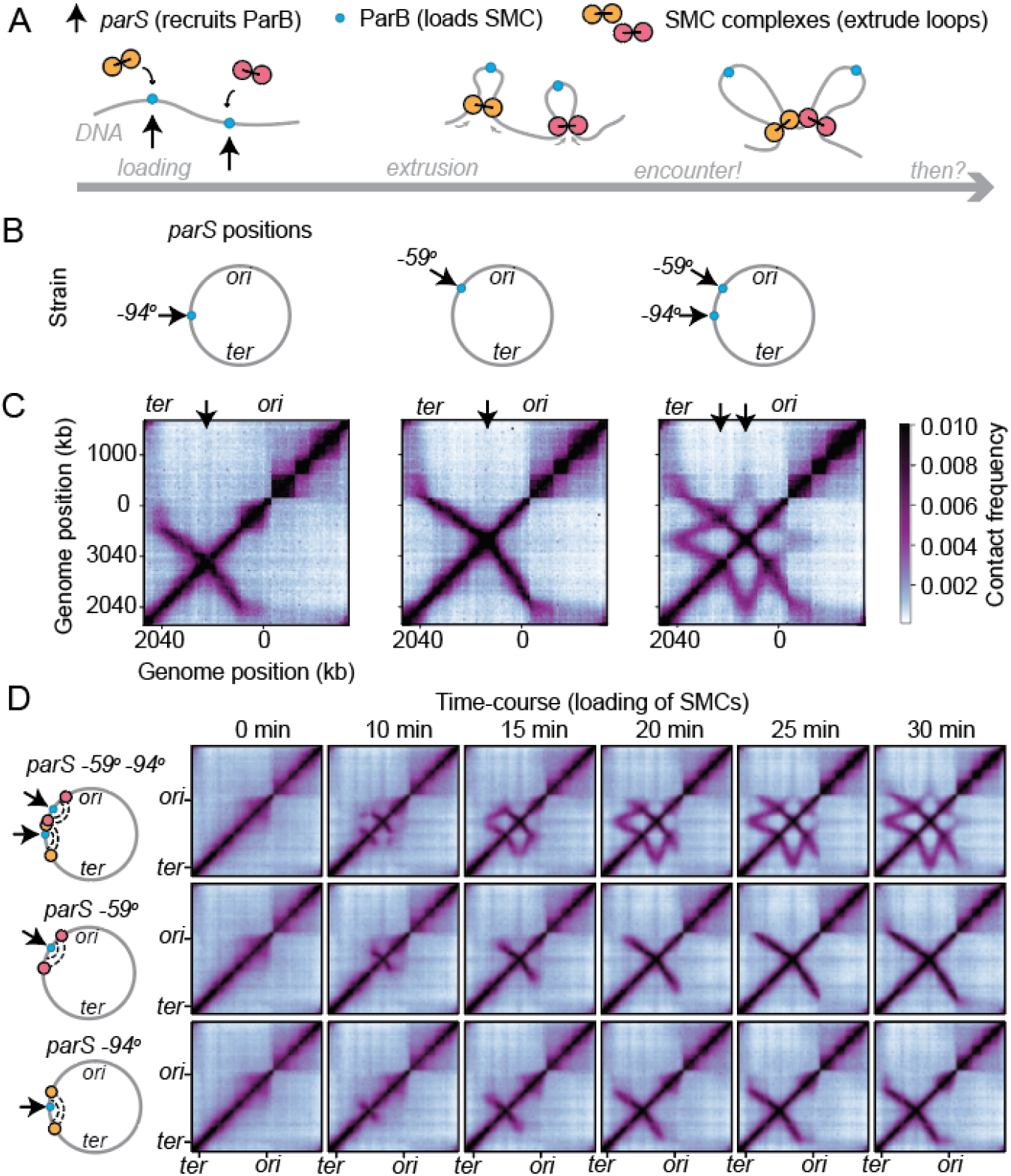
Experimental system to study the effect of “collisions” between SMC complexes. **(A)** Setup of experiment. **(B)** Schematic of strains indicating the inserted positions of single *parS* sites inserted on the chromosome of *B. subtilis*. **(C)** Hi-C performed on G1-arrested cells. Strains contain a single *parS* site at −94° (2981 kb, left), −59° (3377 kb, middle) and at both sites (right). **(D)** Time course experiment following induction of ParB (SMC loading protein) for the indicated times. The schematic illustrates the paths of SMC loop extruders superimposed on the chromosome for each strain at 10 min following ParB induction.

Consistent with our previous findings, strains containing single condensin loading sites at −94° or −59° (i.e. genome positions of 2981 kb or 3377 kb out of 4033 kb, Fig S1A) displayed DNA juxtaposition, or “lines” on the Hi-C map, indicative of large tracks of DNA being brought together in a hairpin-like structure (Fig. 1C, left and center panels) ^3,29^. In striking contrast, a strain with both of these *parS* sites exhibited a complex star-shaped pattern (Fig. 1C, right). This pattern has additional features that are absent from strains with single *parS* sites, indicating that non-trivial interactions occur between condensins translocating from opposing sites. Hi-C performed for the same strains growing in asynchronous cultures revealed similar patterns, albeit less intense, showing that these patterns are not specific to G1-arrest (Fig. S1B). To understand how a star-shaped pattern emerged, we performed a time-course Hi-C experiment in cells with an IPTG-inducible expression of the condensin loader, ParB, as the sole source of the ParB protein (Fig. 1D). We took samples in the absence of IPTG and at 5-minute intervals after its addition. By tracking the juxtaposition of DNA flanking the *parS* site over time, we measured the rates of DNA loop extrusion by condensin. In the strains with a single *parS* site, the extrusion rate was ~0.8 kb/s towards the replication terminus (*ter*) and ~0.6 kb/s towards the replication origin (*ori)*, similar to previous measurements ^3,29^. In contrast, in the strain with both *parS* sites, the extrusion rate in the section between the *parS* sites (i.e. where condensins move towards one another), was lower by a factor of ~1.2, but outside that section the rates remained unaltered (Fig. S2A; see **Supplemental Information**). This slowdown is most evident from the change in the tilt of the lines when comparing to strains with single *parS* sites (Fig. S2B, S2C). These results indicate that condensins translocating towards one another can slow each other down. We thus investigated the underlying mechanism by which condensins interact to create such complex chromosome folding patterns.

We first broke down the star-shaped Hi-C interaction pattern into different line segments and investigated how these lines may be explained by a process of DNA loop extrusion by condensins (Fig. 2A). Lines 1 and 2, similar to those seen in maps for strains with one *parS* site, can be formed by single condensins (making “singlet contacts”) as they translocate away from their respective *parS* loading sites and juxtapose the flanking DNA. In contrast, Lines 3, 4 and 5, are likely formed by interactions between condensins, and we provide below a possible origin for these lines: Line 3 can emerge when two condensins coming from different *parS* sites meet in between the *parS* sites. In addition to each of their singlet contacts (i.e. on Lines 1 and 2), they produce another contact by bridging DNA along their flanks (Fig. 2A ii). Since condensins in different cells can meet at different genome positions, the location of the additional contact varies. Thus, when averaged over a population of cells, the contacts mediated by condensin collisions (i.e. “collision-doublets”) result in Line 3 (Fig. 2A ii, Fig. S3A). Line 4 can emerge if the condensins’ meeting point is at the *parS* site. For example, if one condensin from *parS* site *S2* extrudes past the site *S1*, and a second condensin loads at site *S1* at that moment, then the condensins enter into a “nested-doublet” configuration. As long as the two condensins maintain their contact with each other and continue to extrude DNA, they generate Line 4 over time (Fig. 2A iii). Finally, if the second condensin (loaded from *S1*) of the “nested-doublet” configuration meets a third condensin (loaded from *S2*), then different meeting points between three condensins produces the contacts of Line 5 (see Fig. 2A iv) in addition to Lines 3 and 4. It is possible to envision an alternative mechanism for the formation of Line 5 (and Line 4) (Stephan Gruber, personal communication) (Fig. S4), whereby ParB molecules form a “temporary loading site” at a mirrored *parS1* location on the juxtaposed DNA; however, we can rule this out on theoretical grounds as well as experimentally (see **Supplemental Information**, Fig. S4B). Thus, our line-based decomposition provides a framework for interpreting complex Hi-C patterns as assemblies of condensins and describes a possible series of events leading to each of the lines of the star-shaped pattern.

**Figure 2:**
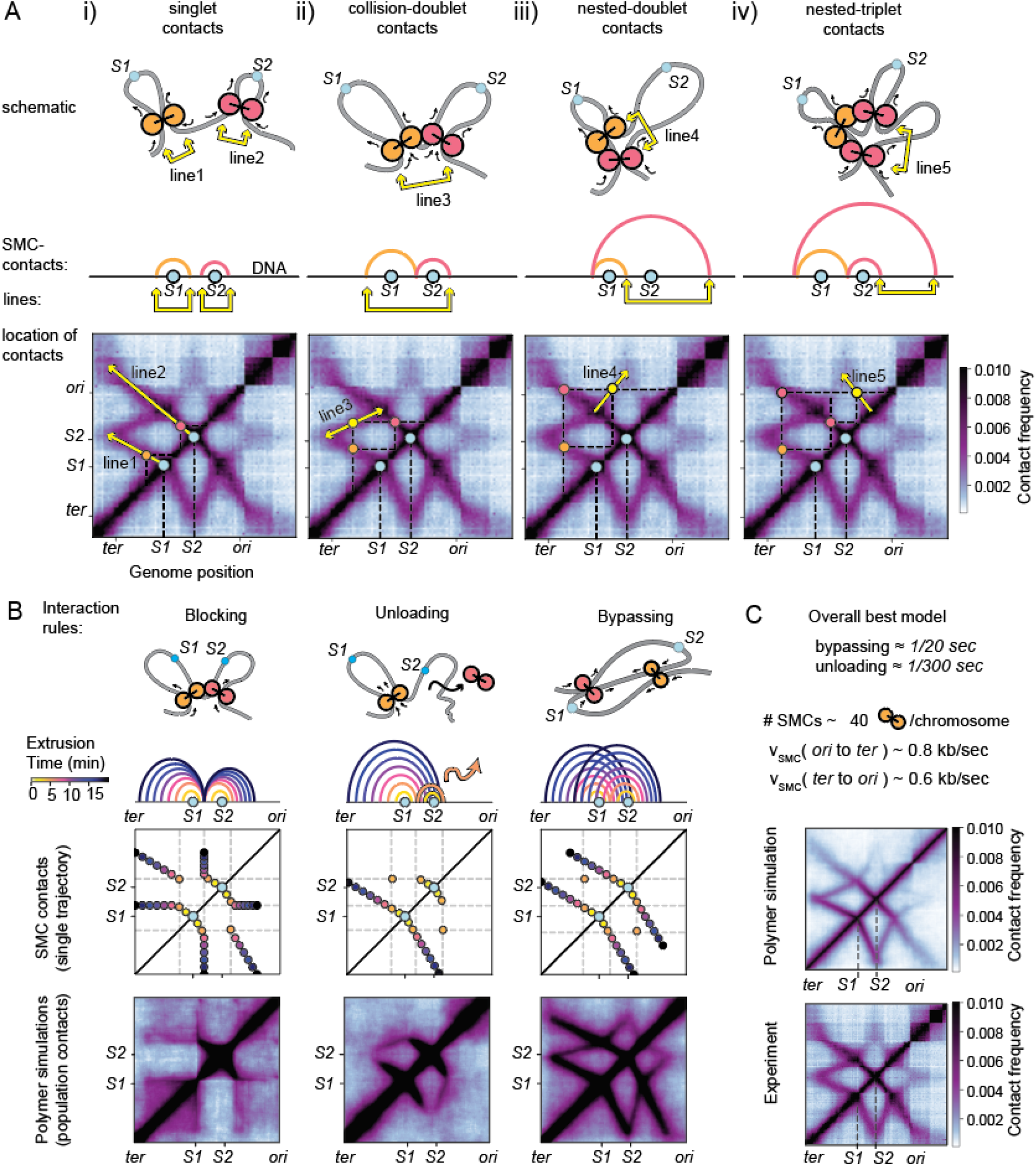
Specific interactions between condensins leave unique Hi-C signatures. **(A)** Diagramatic decomposition of the Hi-C map into assemblies of condensins. The schematic diagram (top row) and the arch-diagram representation (middle row) of the condensin assemblies, are superimposed on a Hi-C map (bottom row). Locations of condensin-mediated contacts are depicted either by a yellow arrow (top, middle), or by a yellow dot (bottom). Condensin loading sites are S1 and S2 (blue dot). Condensins loaded from S1 are colored orange and from S2 pink. These colors are consistent between rows to facilitate comparison. **(B)** Possible interaction rules of condensins (blocking, unloading, bypassing). The schematic diagram (top row) illustrates the interaction, and the arch diagram captures the 1D (second row) and the 2D Hi-C-like contact trace (third row) captures the spatio-temporal behaviour of the interaction of a single illustrative event by a pair of extrusion complexes; extrusion time is shown over a 15 min period, and times are indicated by arch or dot colours. A 3D polymer simulation and the result contact map for each interaction rule is shown on the bottom row. A broader parameter sweep can be found in Figs. S5, S7 and S11. (**C**) A parameter sweep over the three interaction rules, accounting for different rates gives a best-match model. For N=40 extrusion complexes per chromosome, we find that bypassing rates are ~1/20 s^−1^ and unloading rates are ~1/300 s^−1^. A comparison between the experimental data and a 3D polymer simulation of the model is shown.

Next, we used polymer simulations to understand how the patterns observed by Hi-C emerge from the rules of engagement between condensins. In our simulations, each loop extruding factor was represented by two connected motor subunits ^29–31^ (e.g. Fig. 2A). By translocating away from their loading site, the connected motors bring genomic loci into spatial proximity. Based on previous studies of *B. subtilis* condensins ^32^, we allowed loop extruders to load anywhere on the genome, but with a preferential bias (see below) such that most loaded at *parS* sites. Since previous studies showed that two motor activities of the same loop extruding factor are independent of each other, we allowed continued extrusion by a motor subunit even if the other subunit’s translocation was blocked ^3,4,29^. From the simulations of loop extrusion on a 3D polymer, we then created Hi-C-like contact maps ^30,31^(see **Supplemental Information**).

Our attention turned to three main rules of interaction between condensins: blocking, unloading, or bypassing (Fig. 2B, Fig. S3B). We also explored various other models, including 3D interactions between extruders, the effect of sticky DNA, the effect of extruder subunits reversing direction after collision, among others (Fig. S4). However, we ruled out these other models due to their inability to create Lines 4 and 5, or because they generated lines not observed experimentally (Fig. S4).

Similar to prior work, we first considered models involving only blocking ^18,23,30^, and extended the model to include a facilitated unloading of blocked condensins. We allowed collided motor subunits to pause (in the blocked state) before unloading with a specified rate. By sweeping over a broad range of unloading rates and condensin numbers, we found that it was not possible to reproduce Lines 4 and 5 at intensities visible by contact maps (Fig. S5). The failure of this class of models in reproducing Lines 4 and 5 is due to an inability to efficiently form nested configurations: For example, with few condensins per chromosome, it is easy for a condensin loaded at *parS* site *S2* to reach *parS* site *S1*; however, a small loading rate due to few condensins, makes it unlikely that a second condensin will bind to *S1* at the moment the *S2* condensin extrudes past it. With high numbers of condensins, the loading rate at each *parS* site is larger; however, traffic jams due to condensin collisions between the *S1 and S2* sites prevent most condensins from ever reaching the opposing site. Therefore, the blocking and unloading model results in low numbers of nested doublet configurations and cannot create Lines 4 and 5 at the intensities observed experimentally (see theory, **Supplemental Information**).

We thus extended the blocking-only model by allowing condensins to bypass one another, which was also motivated by recent single molecule experiments ^12^. In this blocking and bypassing model, we assumed that, when two condensins meet, the collided subunits pause but can traverse each other with some specified rate; however, we did not allow facilitated unloading. Strikingly, the blocking and bypassing model was sufficient to robustly reproduce the star-shaped Hi-C pattern (Figs. 2B, S6A). Moreover, bypassing produced the observed “tilting” of Lines 1 and 2 away from each other for certain bypassing rate and condensin number combinations (Fig. S6B). Nevertheless, the bypassing mechanism by itself generated Lines 4 and 5 more intensely than observed experimentally, suggesting that too many SMC complexes were entering nested configurations (Fig. 2A). Therefore, we added back the facilitated unloading assumption to the blocking and bypassing model. This allowed us to tune relative intensities of Lines 3, 4 and 5 and obtain Hi-C maps that looked strikingly similar to the experimental data (Fig. S7; see **Supplemental Information**). Of all the models that were tested, the combined bypassing and unloading model was the only one that produced all the Lines 1-5 at the same time, with the observed relative intensities.

The resulting integrated model included the rules for condensin encounters (i.e. bypassing and facilitated unloading), as well as the rules for basal condensin dynamics and totaled six parameters: the bypassing rate, the facilitated unloading rate, the number of condensins per chromosome, the condensin loading rates at *parS* sites versus other sites, and the spontaneous dissociation rate of condensins in the absence of collisions. Uniquely, we found that we could fix all the six model parameters experimentally using a combination of Hi-C, ChIP-seq data and theoretical constraints between parameters, finding a unique region of parameter space which best fit all of the available data (Figs. S8, S9, S10, S11; see **Supplemental Information**).

In the best models, there were 25-45 condensins present on each chromosome after 1 hour of G1-arrest. Condensins paused for ~20 s when they met each other before either bypassing or unloading from the chromosome (Fig 2C). Moreover, we found that bypassing was ~10-20 times more likely to occur than unloading (i.e. the bypassing rate was ~0.03-0.1 s^−1^ whereas unloading was ~0.002-0.005 s^−1^). Thus, the bypassing mode of conflict resolution dominated over unloading and was essential for explaining the observed lines on the Hi-C map mediated by SMC complex dynamics. Fixing the rates for bypassing at 0.05 s^−1^ and unloading at 0.003 s^−1^, and with 40 extruders/chromosome, polymer simulations quantitatively reproduced the condensin-mediated Lines 1-5 seen in the experimental Hi-C data (Fig. 2C), and raised the possibility of predicting chromosome folding in other engineered strains.

Thus, we investigated whether the bypassing and unloading rules were generally applicable to condensin encounters. We generated seven other strains containing two *parS* sites at various locations and two strains containing three *parS* sites and performed Hi-C in G1-arrested cells (Fig. 3A, 3B) and exponentially growing cells (Fig. S12). These engineered strains produced an impressive diversity of Hi-C contacts patterns, which depended on the relative spacing and positioning of condensin loading sites. Nevertheless, all the complex multi-layered interactions features could be understood via the descriptions of condensin-mediated contacts (Fig. 2A). Strikingly, using the same parameter values as above (i.e. from Fig. 2C), the model of condensin bypassing and unloading accurately reproduced all the emergent Hi-C contact features, showing strong agreement with all nine strains (Fig. 3A, 3B).

**Figure 3:**
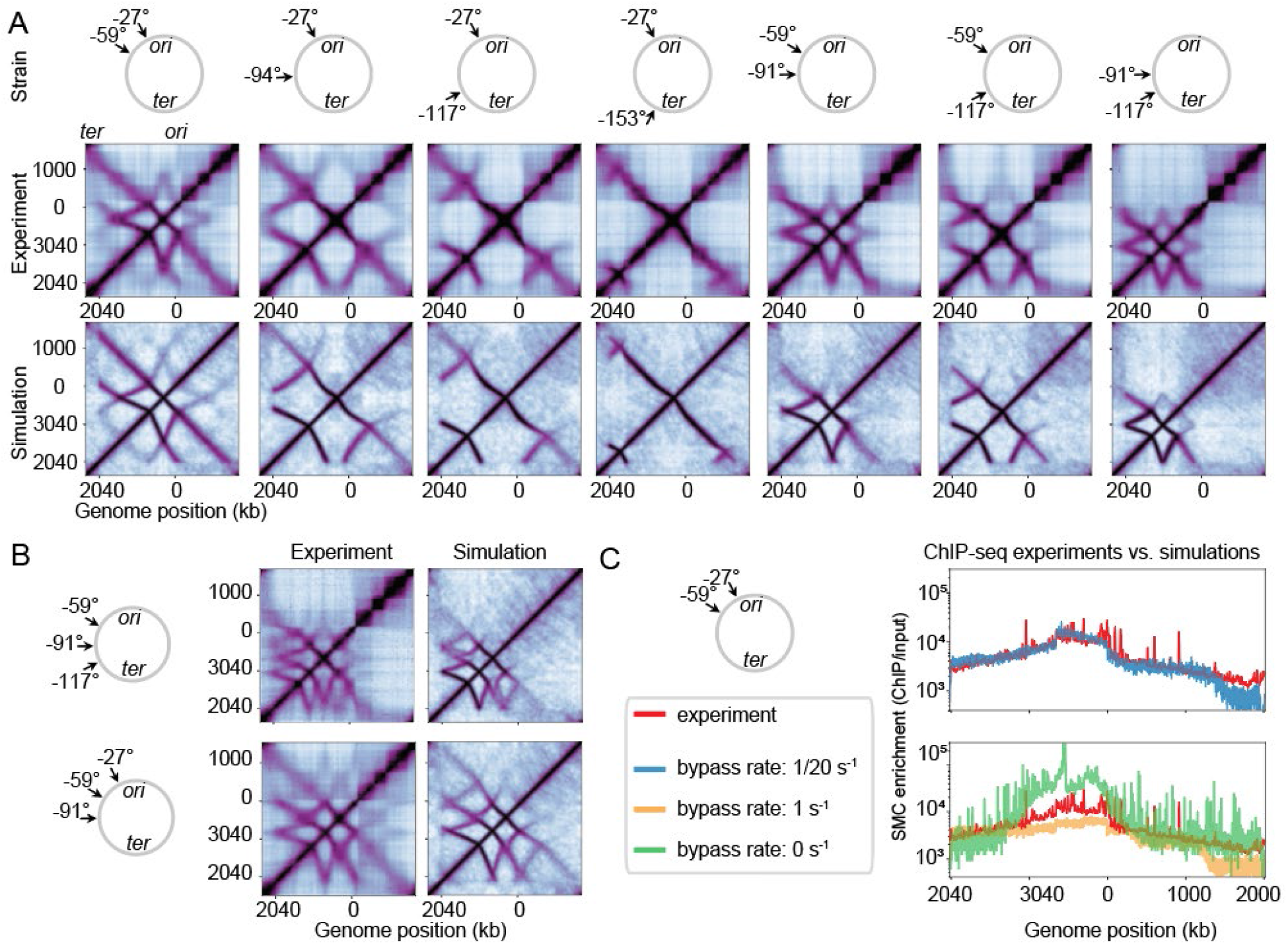
Validating and testing a model of *in vivo* Z-loop formation. Data and simulations of Hi-C maps of strains containing two *parS* sites **(A)** or three *parS* sites **(B)**; all models use the same parameters as identified in Fig. 2C. **(C)** A comparison of condensin occupancy as assayed experimentally by ChIP-seq and derived from our model for different rates of bypassing; notably, a bypassing rate of ~1/20 s^−1^ identifies the best-fit model, independently of the Hi-C. Our models work well to capture the genome-wide trends of condensin occupancy except near the terminus region, indicating that our understanding of the condensin loading at *parS* sites and interaction rules are good, but future work needs to be done to elucidate how condensins interact with the terminus.

As an independent way of investigating the consequences of condensin encounters, we determined condensin distributions by performing SMC ChIP-seq and compared experiments to our model predictions (Fig. 3C). We found that a bypassing rate of ~1/20 s^−1^ was necessary for quantitative agreement between experiments and simulations (Fig. 3C). With the bypassing rate too low, condensins tended to accumulate strongly near the loading sites; with the bypassing rate too high, the occupancy profile was too flat. This rate of bypassing (~1/20 s^−1^) obtained by modelling of ChIP-seq is in strong agreement with the value inferred from modelling of Hi-C data. Together, the agreement of the model with both Hi-C and ChIP-seq lends strong support for the notion that condensins can translocate past one another on DNA *in vivo* after short pauses.

Having studied the effects of changing loading site positions and spacings, we next studied the effect of time on chromosome reorganization after G1-arrest. In wild-type cells, which harbour nine *parS* sites, the spatial chromosome organization changes dramatically; the most prominent feature of the Hi-C map (the central diagonal) vanishes after two hours and is replaced by two smaller tilted lines ^25^. However, before examining the time-dependent changes in the wild-type system, we investigated the time course in a simplified system with one and two *parS* sites. Over a two-hour window, although no changes occurred to the Hi-C Lines 1 or 2 with one *parS* site (Fig. S13A), we observed major changes in strains with two sites (Fig. 4A, S13B). Specifically, the star-shape became progressively larger due to an increased tilt of Lines 1 and 2 away from each other (Fig. 4A, S13B). This indicated that the condensin translocation between the *parS* sites was further slowed down over time and demonstrated that the observed changes were due to interactions of condensins translocating from different *parS* sites. By simulations, we could also achieve a similar effect (Fig. 4A, S6). By increasing the numbers of loop extruders present on the chromosome we obtained more frequent condensin collisions which led to an overall slowing down of extrusion between *parS* sites and the larger star-shaped pattern. The numbers of loop extruders per chromosome necessary to recapitulate the Hi-C data were 40±10, 60±15, 90±20 (Fig. 4A). Reassuringly, the numbers of extruders per chromosome that gave the best agreement with Hi-C also independently reproduced SMC ChIP-seq profiles for the specific time points (Fig. 4B). We thus hypothesized that continued protein synthesis after replication inhibition resulted in a higher number of condensins per DNA molecule.

**Figure 4:**
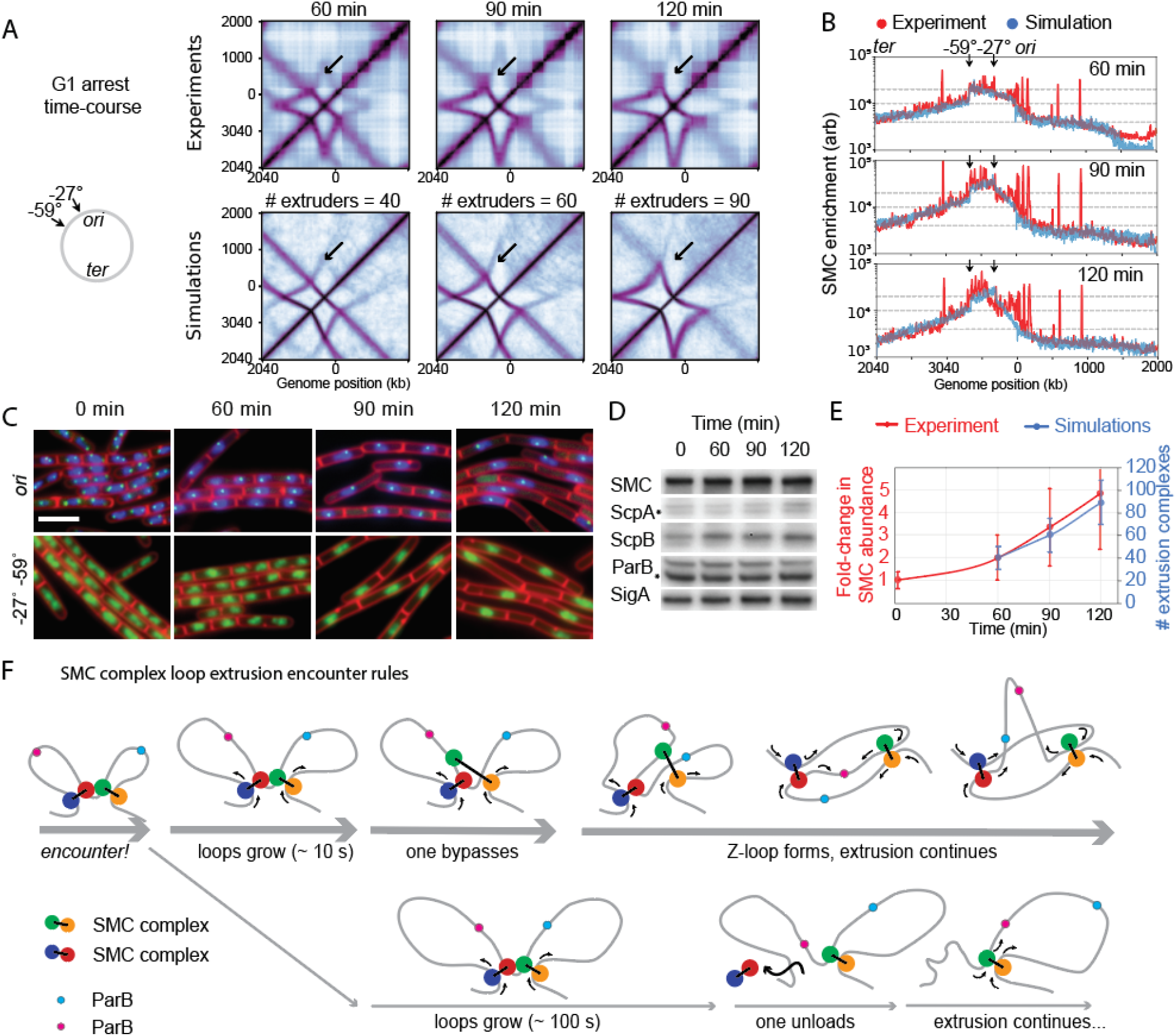
Numbers of SMC complexes per chromosome tunes the shape of contact maps. A time-course of Hi-C **(A)** and SMC ChIP-seq (**B**) tracking chromosome structure changes and condensin occupancy following replication arrest by SirA; experiments are compared to our simulations where the numbers of condensins per chromosome are increased but all simulation parameters are otherwise kept the same as in Fig. 2C. **(C)** Fluorescence microscopy of tagged chromosome loci (green) marking the *ori* (top row), and the *−27*° *−59*° positions (bottom row); the nucleoid is shown in blue, and a cell wall marker is in red. Notably, the cells increase in length, and *ori* counts per nucleoid become one. **(D)** Quantitative western blot for various protein components of the SMC complex; results indicate that after replication arrest the density of SMC complex components per cell remains largely unaltered. **(E)** Quantification of the number of extrusion complexes per cell (assuming extrusion complexes as dimers of condensins); numbers are calculated using the fluorescence microscopy, Western blot, and whole-genome sequencing data quantifications; notably, the data are in good agreement with the Hi-C derived numbers obtained via simulations and shows that condensin numbers/chromosome increases after replication inhibition. **(F)** Final model schematic illustrating how SMC complex encounters are resolved. Notably, upon encounter, SMC complexes first mutually block one another and then may resolve the conflict by either bypassing (top row), or unloading from the DNA (bottom row). The bypassing mode of conflict resolution occurs at least 10 times more frequently than unloading (as indicated by the thickness of the arrows).

To quantify the change in condensin abundance experimentally, we measured the chromosome copy numbers per cell and condensin protein abundances over time: Marker frequency analyses^33^ by whole genome sequencing and fluorescence microscopy showed that cells retained only one copy of the genome per cell for the duration of the experiment (Fig. 4C, S9). Immunoblot analyses of cells growing in the same conditions showed that ParB and condensin subunit levels per unit cell mass remained constant over time (Fig. 4D). However, we found that at 90 min and 120 min after G1 arrest, the nucleated cells’ length/mass increased to 1.7-fold and 2.4-fold of the 60 min value (Fig 4C, S14). From the increased cell lengths and constant density of condensins, we inferred the relative changes in condensin numbers per chromosome (Fig. 4E). These fold-change values in SMC complex numbers are in good agreement with the numbers of loop extruders independently identified by Hi-C and simulations above (Fig. 4E). Thus, continued protein synthesis after replication inhibition leads to increased numbers of condensins per chromosome.

Next, we directly tested the role of condensin abundance on chromosomal organization by perturbation. We hypothesized that condensin overexpression would lead to a faster evolution of the observed Hi-C patterns in a G1 arrest time course. Consistently, we observed this trend experimentally in a strain with two *parS* sites (Fig. S15): at the 60 min mark, we saw the traces typical of 90 min in the absence of condensin overexpression. This confirms the role of condensin abundance in tuning chromosome spatial organization and the changing shapes of the Hi-C interaction patterns.

Finally, we studied the most complex systems: strains with three *parS* sites or 9 *parS* sites (i.e. wild-type cells), and investigated the mystery of the vanishing “central” lines. In these strains, the disappearance of the central line in Hi-C was accompanied by the accumulation of condensins between the *parS* sites by SMC ChIP-seq (Fig. S16, S17). Despite the complexity of the changes over time, our model captured these effects (Fig. S16, S17) and helped to understand what was happening: In normal growth conditions, with basal condensin levels, collisions between condensins from adjacent *parS* sites are resolved by bypassing (i.e. in ~20 s) before the next extrusion complex arrives. However, the increased number of condensins makes the rate of new collisions higher than the rate of bypassing. This effect is particularly strong for the central sites, where extruders are jammed in from both sides. This finally results in effective extrusion only from the out-most *parS* sites and gives rise to the disappearance of the central line (Fig. S16, S17). We conclude that bypassing plays an important role in preventing traffic jams between condensins in wild-type cells under normal growth conditions, by allowing productive extrusion from multiple neighboring *parS* sites.

Thus, a model where condensins can traverse one another on the chromosome after momentary pausing is consistent with results from many strains and conditions tested here (Fig. 4F). Our study demonstrates that by harnessing the condensin-ParB-*parS* system, we can create complex chromosome folding patterns not seen before in natural systems, which helped understand what is happening in the wild-type cells. Strikingly, these structures could be predicted by a quantitative model of condensin dynamics, which was central to identifying the bypassing mechanism as a key feature of *B. subtilis* condensin loop extrusion. We inferred that condensins can traverse past each other within ~20 s of an encounter *in vivo*. This time scale is consistent with the *in vitro* times of ~8 s measured by single molecule experiments for yeast condensins to traverse past one another on naked DNA ^12^. These times are also consistent with the ~10 s in *B. subtilis* to traverse sites of active transcription as shown previously ^29^. Together, these results suggest that the phenomenon of condensins traversing past one another, and other steric obstacles, may be general to many species and processes.

In specific situations, we found it is possible to overwhelm the bypassing mechanism and create condensin traffic jams. The jamming, caused by elevated numbers of chromosome-bound condensins, is similar to the phenomenon where high RNA polymerase traffic (that opposes the direction of condensin translocation, e.g. at rRNA genes) leads to the accumulation (and pausing) of loop extruders at transcription end sites ^4,29,34^. At first glance, condensin molecules bypassing each other to form structures such as z-loops appear to tangle the DNA. However, bypassing generally helps avoid traffic jams formed with condensins loaded at adjacent sites. This is important in bacteria since *parS* sites often occur in multiple copies close to the *ori* ^20^. As an example, if bypassing were not a feature of condensins in wild-type *B. subtilis* cells, then pervasive tethers between the *ori* and other genome positions would frequently occur, potentially affecting *ori* segregation (Fig. S18). To prevent such long-range tethers, *B. subtilis* cells would have to organize chromosomes with no more than 1 to 4 condensins per chromosome (i.e. <20% of experimentally measured values ^32^). In such a case, however, the chromosome arm juxtaposition is poor as seen by simulations (Fig. S18), and much weaker than seen experimentally ^25^. Thus, bypassing is an essential property that allows multiple *parS* sites to function together efficiently, not only in engineered strains, but also in wild-type cells in exponential growth conditions.

In eukaryotes, bypassing can help promote chromosome compaction and sister chromatid segregation ^30,35^. However, we hypothesize that bypassing could have a function beyond compaction and segregation. For instance, bypassing of obstacles and other SMC complexes could potentially facilitate spreading of cis-related chromatin marks (e.g. around a DNA double-strand break ^36–38^), or help trafficking of various factors along the chromosome ^39,40^. Speculatively, if the ability to bypass obstacles is rampant, cells may have developed specific mechanisms to control this process and stop extrusion (e.g. CTCFs for cohesins ^41^).

A future challenge of the field is to investigate the molecular mechanism of bypassing using biochemical and structural approaches. In addition, single molecule approaches will be powerful to determine the ability and efficiency of various SMC complexes to bypass one another, as shown previously ^12^. However, the targeted SMC complex loading approach (as we showed here) can produce distinct signatures visible by Hi-C that can help distinguish bypassing from other mechanisms. Employing this idea in a eukaryotic system could be very powerful to investigate if bypassing occurs *in vivo* in eukaryotes.

In summary, we have shown that *B. subtilis* condensin SMC complexes can resolve encounters by simply translocating past one another allowing them to spatially organize a functional and busy genome.

## Supporting information

Supplemental Calculations

## Author contributions

H.B.B. and X.W. conceived of the project; H.B.B. developed the theory and models and analyzed data with L.A.M, with input from X.W.; Z.R. and X.W. performed Hi-C experiments and analyses; X.K. and X.W. constructed strains, performed and analyzed ChIP-seq, whole genome sequencing, fluorescence microscopy and immunoblot experiments. H.B.B., L.A.M and X.W. interpreted results and wrote the article with input from all authors.

## Acknowledgements

Hi-C and ChIP-seq data were deposited in the Gene Expression Omnibus (accession no. GSE). We thank Maxim Imakaev and Anton Golobordko for the development of the polychrom simulation package, as well as Edward Banigan, Aafke van den Berg, Kirill Polovnikov and Eli Schantz for discussions. We are grateful to Stephan Gruber, Chase Broedersz, Job Dekker, and Janni Harju for critically evaluating our manuscript and exchanging ideas. We thank Mira Suiter for strain building, Indiana University Center for Genomics and Bioinformatics for assistance with high throughput sequencing, Alan Grossman for anti-SMC and anti-ParB antibodies, David Rudner for anti-ScpA, anti-ScpB and anti-SigA antibodies. Support for this work comes from Startup funds from Indiana University (X.W.), the National Institute of Health Grants U01CA200147 and GM114190 (L.A.M.), H.B.B. was partially supported by a Natural Sciences and Engineering Research Council of Canada Post-Graduate Fellowship (Doctoral). L.A.M. and H.B.B. acknowledge support from the National Institutes of Health Common Fund 4D Nucleome Program (DK107980).

## Supplemental figures

**Figure S1:**
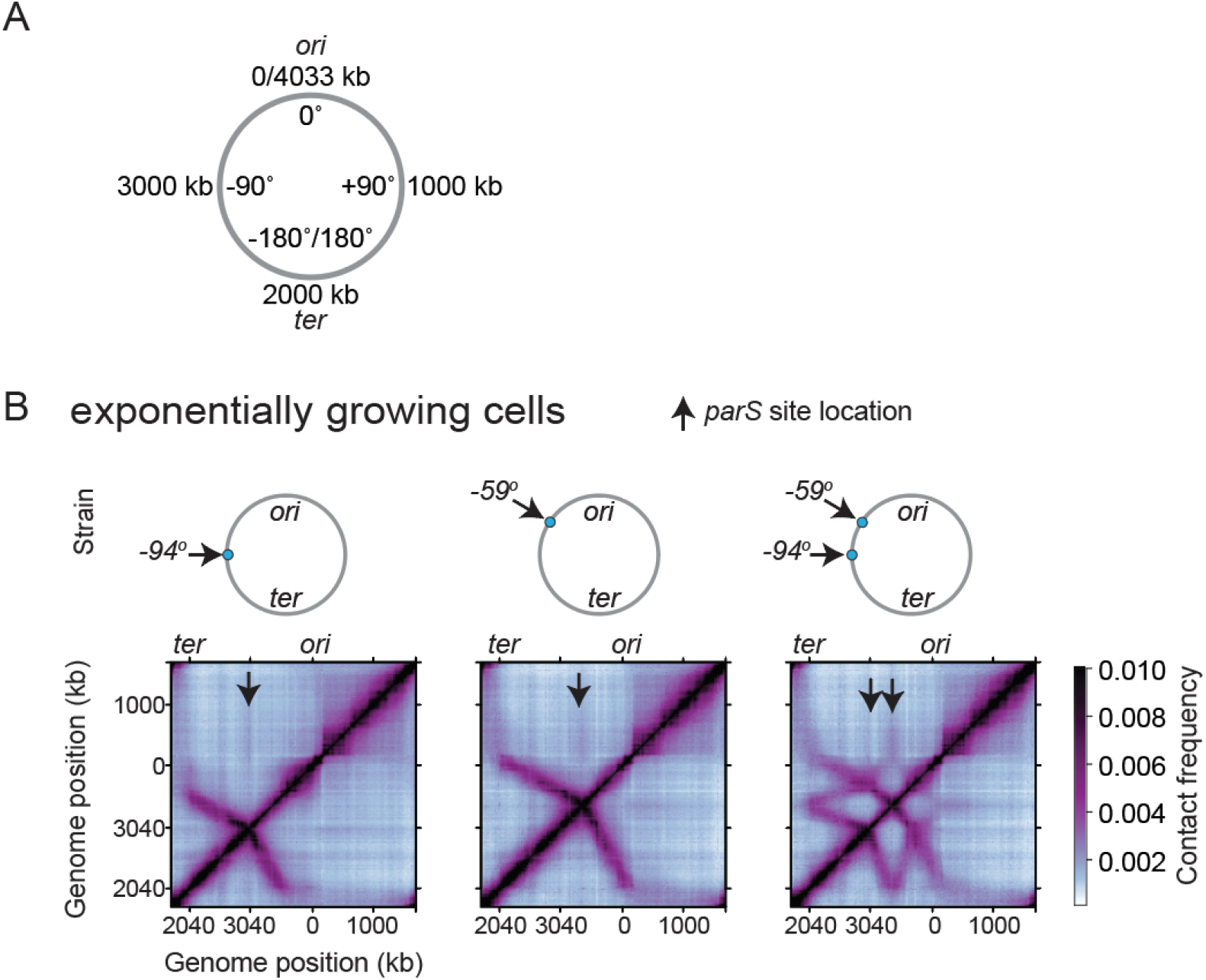
Exponential growth does not strongly affect the Hi-C contact patterns. **(A)** The *B. subtilis* genome is displayed in genomic coordinates (kilobases) and the angular coordinates used to designate the locations of the *parS* sites. (B) Hi-C maps for cells in exponential growth for the strains with *parS* sites at −94°, −59°, and both −94° and −59°. For details on strain names refer to Table S1.

**Figure S2:**
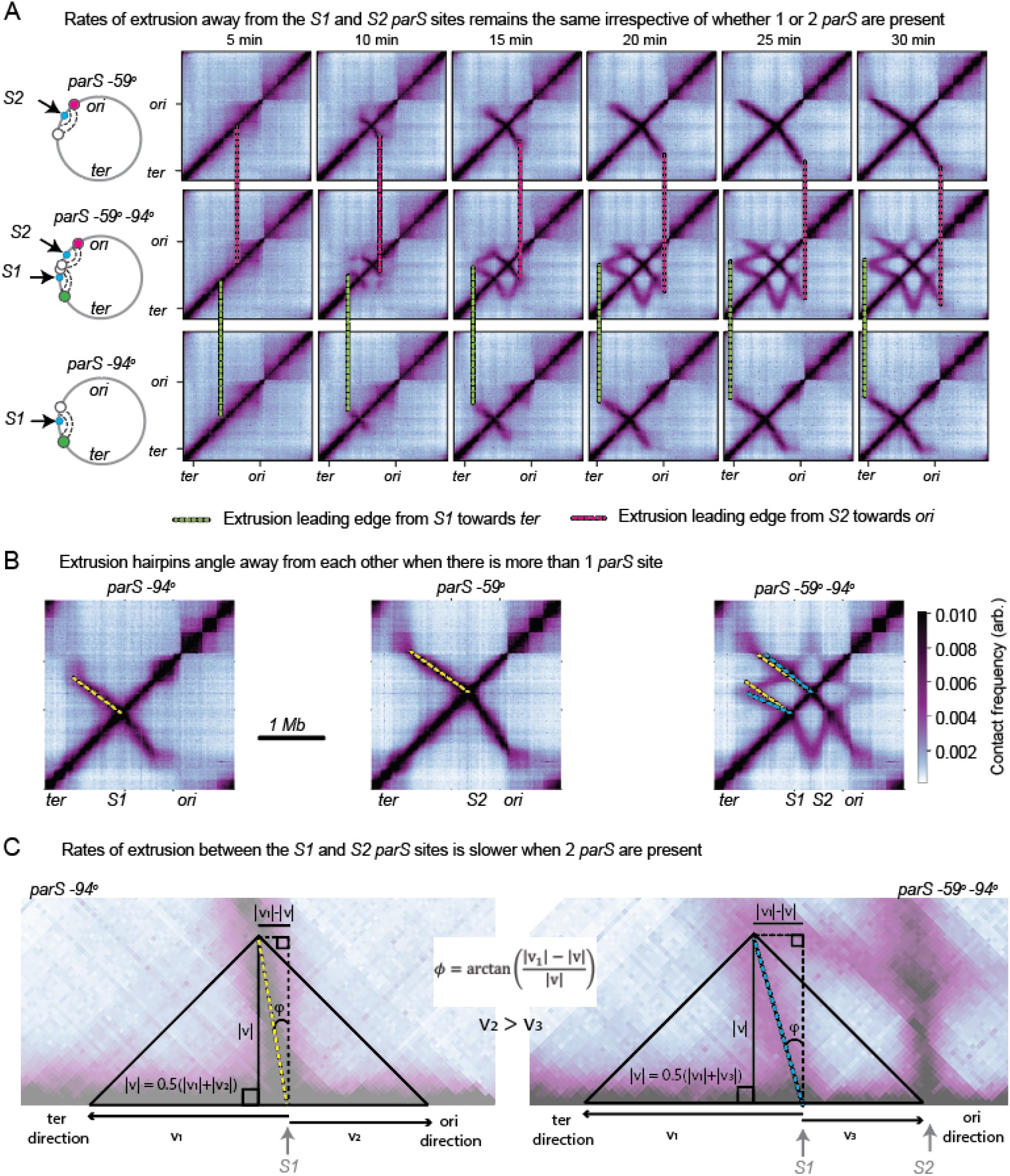
Condensins can slow each other down. **(A)** Time-course (similar to Fig. 1D) comparing the extrusion rates away from (but not between) the S1 and S2 *parS* sites. The green dashed line tracks the leading edge of the hairpin trace as it emerges from the −94° *parS* (S1) site and moves towards the *ter*; the red dashed line tracks the leading edge of the hairpin trace as it emerges from the −59° *parS* (*S2*) site and moves towards the *ori*. **(B)** Demonstration that when two *parS* sites are in one strain, the angle of the hairpin traces changes compared to single *parS* sites. The yellow and blue dashed lines are superimposed on the Hi-C map to help visualize the angle change. **(C)** The relationship between the tilt of the hairpin trace and the loop extrusion speeds (v_1_, v_2_ and v_3_) is captured by a simple geometric relation. The equation shows that for equal v_1_ across strains with one or two *parS* sites (as indicated in panel (**A**)), it follows that v_2_ > v_3_.

**Figure S3:**
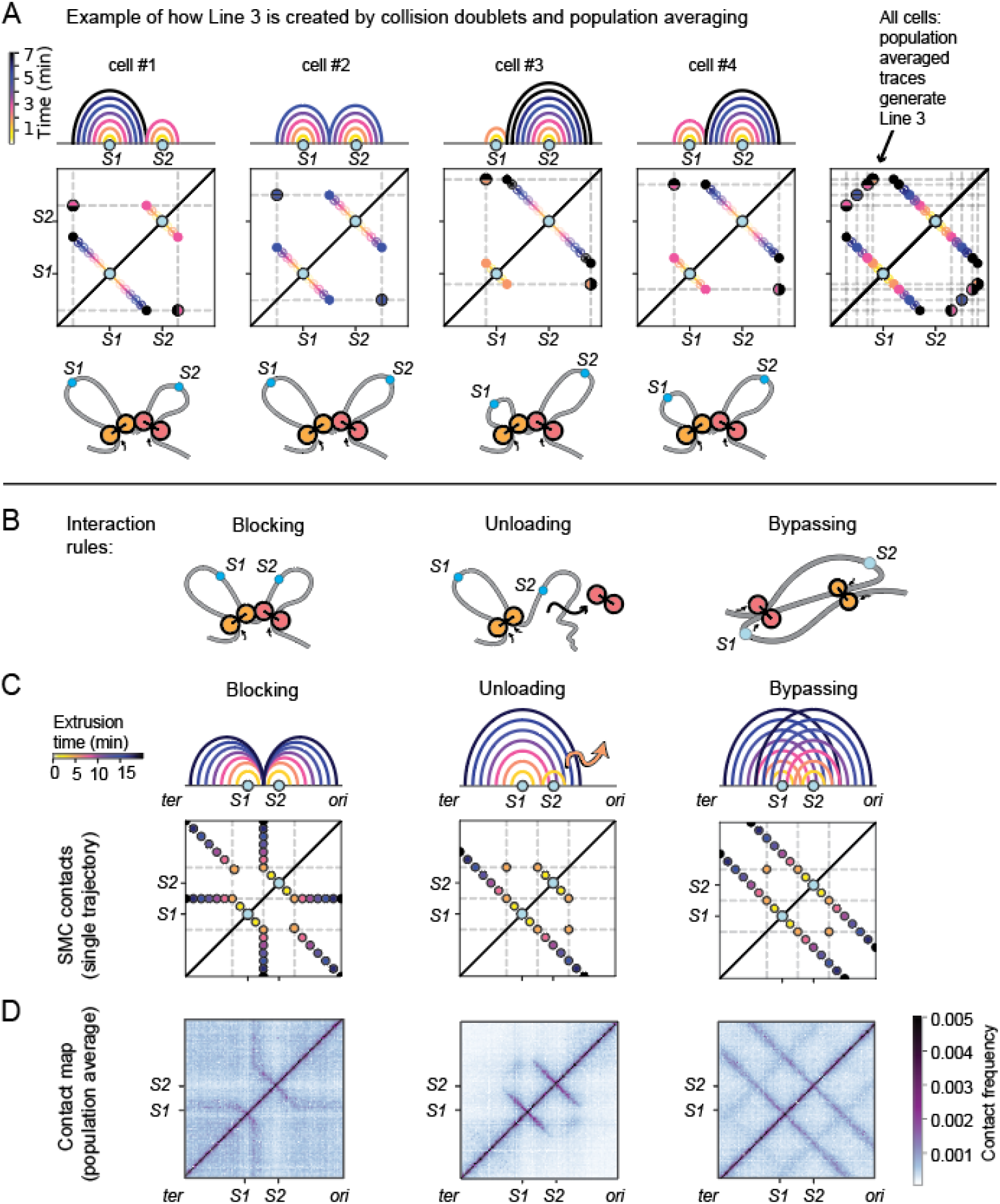
Intuition building for interaction rules (case of equal rates). **(A)** Example of how collision doublet conformations form Line 3. *Top row*: arch diagrams showing the time-course of SMC complexes extruding DNA from their *parS* sites (S1 and S2) up to the point of collision. *Middle row*: 2D traces of the extrusion trajectories; each dot shows the genomic coordinates bridged by a loop extruder at various where the time is indicated the colour. *Bottom row*: schematic showing the loops formed at the point of collision. **(B)** Schematic of the interaction rules for the case where loop extruding factors extrude with equal rates away from their loading sites. **(C)** Arch diagrams corresponding to the genomic positions bridged by single loop extruders over time; *top row*: colours of each arch correspond to a specific time after loading of the extruder on the genome; *bottom row*: 2D representation of the trajectories. **(D)** Hi-C like contact map resulting from a population average over many extruder trajectories; this illustrative map was generated from a loop extrusion simulation coupled to the semi-analytical approach to generating contacts (see Supplemental Information), with N=20 loop extruders on the genome.

**Figure S4:**
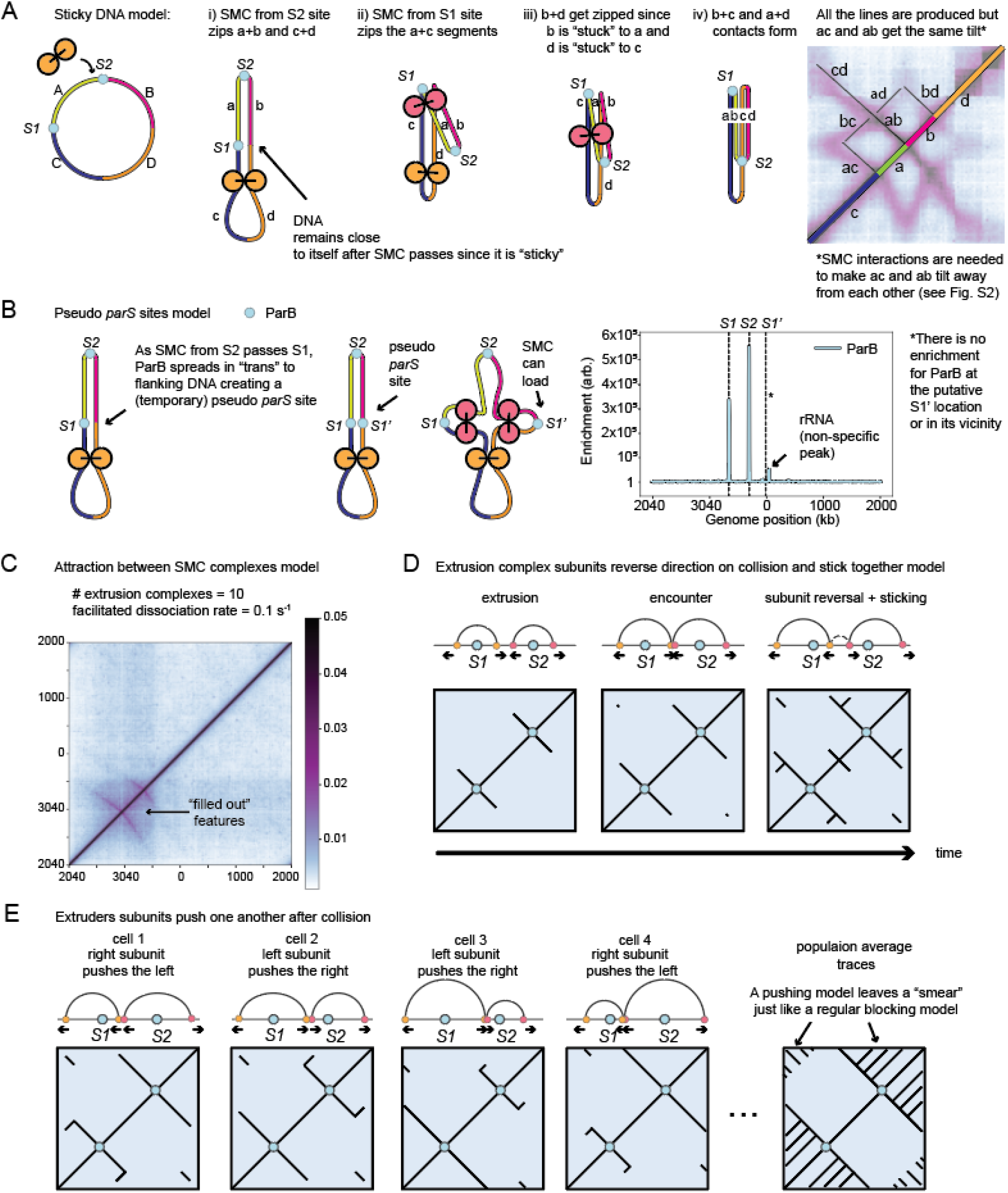
Alternative models we rule out. **(A)** Sticky DNA model. DNA segments (a and b; c and d) stay together after SMC complexes loaded at *S2* pass by. Then when a new SMC complex loads at *S1*, as it generates interactions between segments a and c, it also generates interactions between a and d, b and c because ab and cd are stuck together. Although this could fold the chromosome into a star-shaped pattern and can generate all the lines observed in the Hi-C map, this model does not produce the necessary and observed “tilting” of the lines (e.g. ac and ab) away from each other. The tilting of Hi-C traces in strains with multiple *parS* sites relative to strains with single *parS* sites indicate non-trivial interactions between condensin complexes (leading to slowing down of SMC translocation). **(B)** Pseudo-*parS* sites model. When SMC complexes loaded from *S2* pass *S1*, ParB at *S1* spreads onto the mirror chromosome arm at *S1*’ creating a temporary/pseudo loading site for SMC complexes. This model would predict some accumulation of ParB at the pseudo-*parS* (S1’) site. However, in ChIP-seq experiments, ParB accumulation at the *S1*’ site is not visible for a strain with *parS* at −27° and −59°. We note that the small non-specific ChIP-seq peak to the right of the *S1*’ site (black arrow) at a cluster of rRNA genes which appears irrespective of which antibodies are used (Wang et al, 2017). **(C)** 3-D attraction between SMC complexes. If SMC complexes translocate on different DNA segments are randomly attracted by 3D attractions to each other, the emergent patterns do not resemble a complete star-shape (e.g. Fig. S5), and features emerge as smears as opposed to lines “hollow”. The shown map corresponds to N=10 extruders and k_u_=0.1 s^−1^ and was computed with the semi-analytical model; the strong attraction was created by adding extra random harmonic bonds between half of the extruders. **(D)** Reversal and sticking upon collision. When two DNA extruders meet, they stick to each other and both the inner subunits of the extrusion complex reverse direction. The sticking interaction generates new interactions depicted as a “dashed” line between the orange and magenta subunits in the top, rightmost panel. Bottom panels depict the time-averaged 2D representation of the trajectories. This model produces lines on the interaction maps that are not seen experimentally. **(E)** Subunit pushing upon collision. When two DNA extruders meet, one subunit dominates the other and pushes the other back. This model produces lines on the interaction maps that are not seen experimentally.

**Figure S5:**
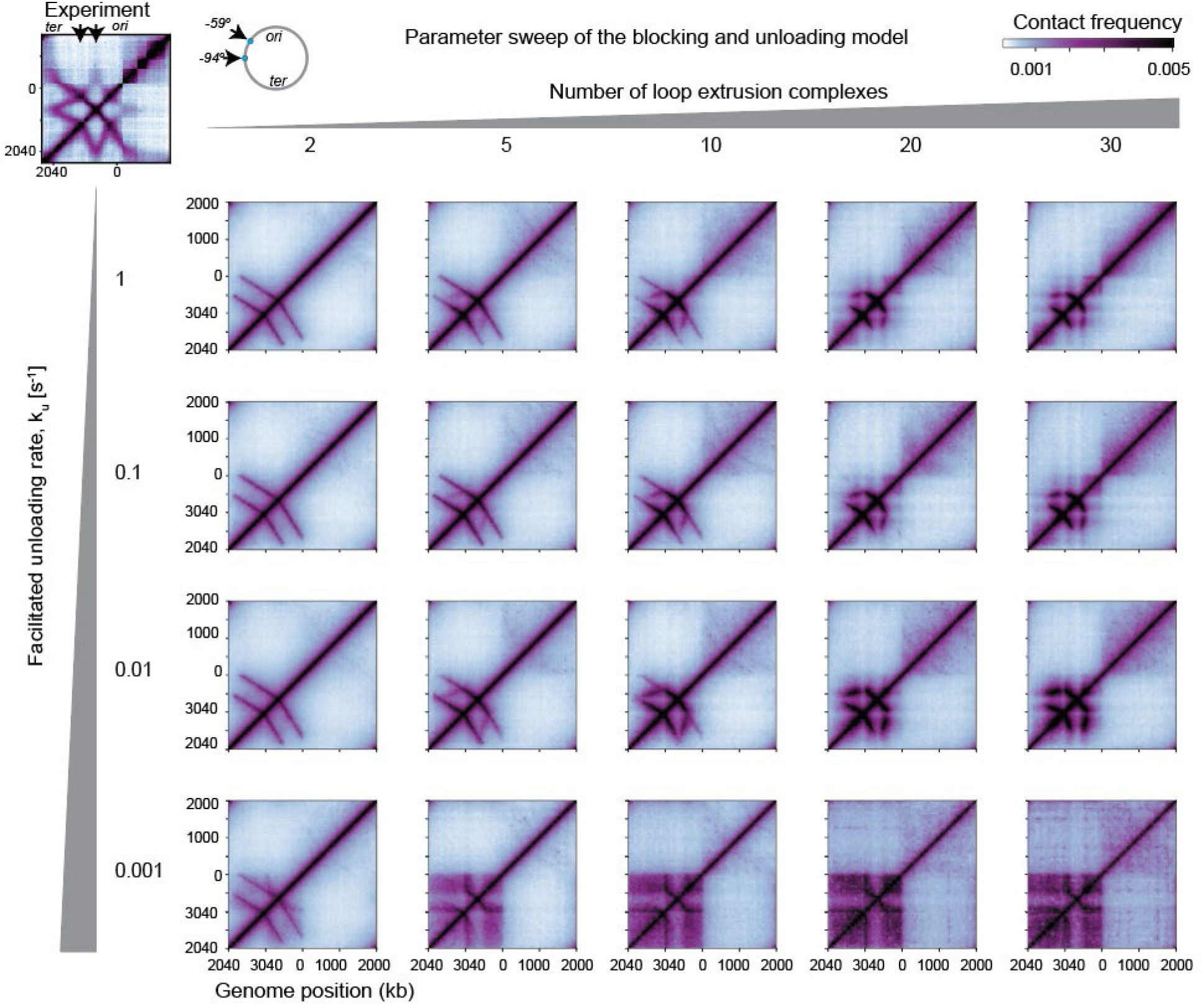
The blocking and unloading model of loop extruder interactions does not produce all the features seen in the Hi-C map. The parameter sweep was conducted for varying numbers of extruders and facilitated dissociation rates. The experimental data (for the strain with *parS* sites at both - 94° and −59°) is shown on the top left of the figure. These contact maps were generated with the semi-analytical approach without making the shortest path approximation as described in Banigan et al, 2020, Appendix 3 (also, see **Supplemental Information**). Notably, Lines 4 and 5 are missing in all of the plots with the blocking and unloading model - this is due to either traffic jams forming between extruders (for high numbers of extruders), or an insufficient loading rate (for low numbers of extruders) preventing the efficient formation of nested doublets and triplets.

**Figure S6:**
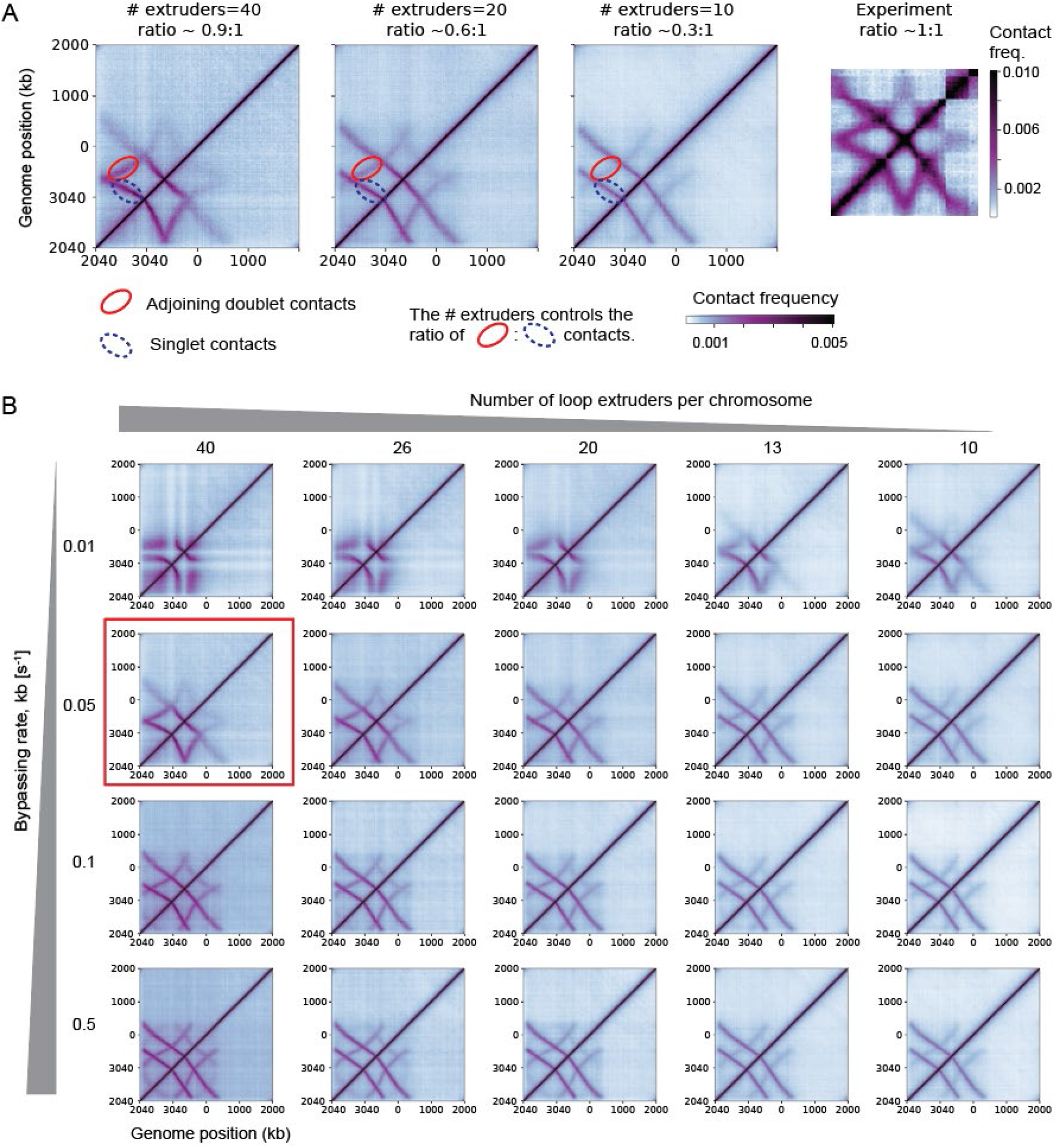
The number of extrusion complexes tunes the relative frequency of singlet-to adjoining-doublet interactions. **(A)** In the experimental Hi-C map for the −59° −94° strain after 60 min of SirA treatment, the frequency of singlet contacts to adjoining doublet contacts is close to 1:1. This is only achieved when the number of condensin extrusion complexes is >40 and for sufficiently high bypassing rates. **(B)** A parameter sweep over the number of extruders and the bypassing rate. From this approach, we obtain a lower bound on the number of extrusion complexes required to produce the experimental Hi-C pattern (i.e. 40 extruders) The best matched parameter combination is shown boxed in red.

**Figure S7:**
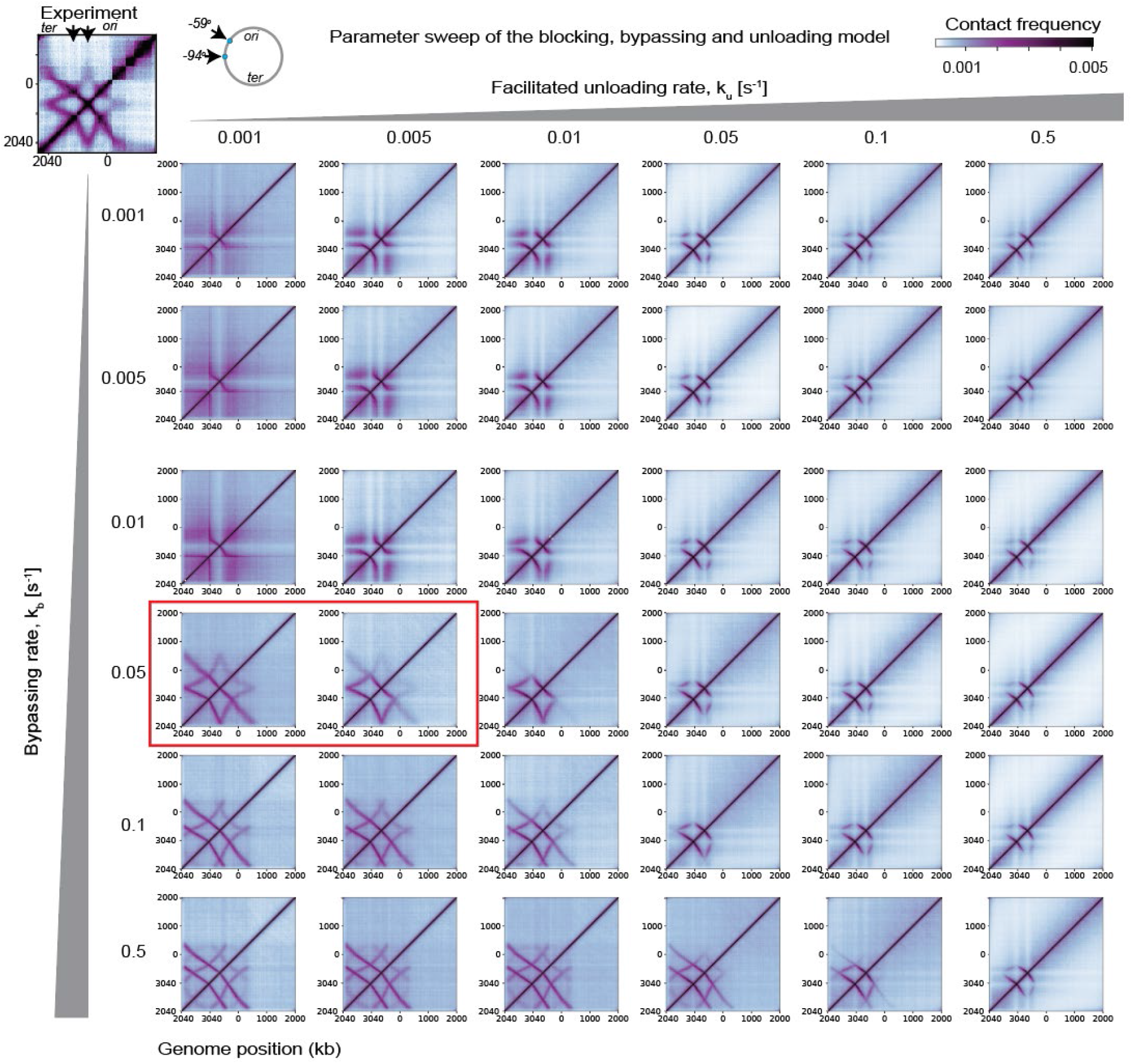
Parameter sweep of the bypassing and unloading rates for N=40 extruders/chromosome. The experimental data (for the strain with *parS* sites at both −94° and −59°) is shown on the top left of the figure, and the model parameter sweep is below. The model with the most similar pattern in both angles of the Hi-C traces and the relative intensities of the different lines corresponds to a bypassing rate of 0.05 s^−1^ and an unloading rate between 0.001 s^−1^ and 0.005 s^−1^. From the sweep, we find that the bypassing rate can control the angle between Hi-C map hairpin structures, while the ratio of the bypassing to unloading rates tunes the relative frequency of nested-doublet and nested-triplet interactions. These contact maps were generated with the semi-analytical approach (see **Supplemental Information**).

**Figure S8:**
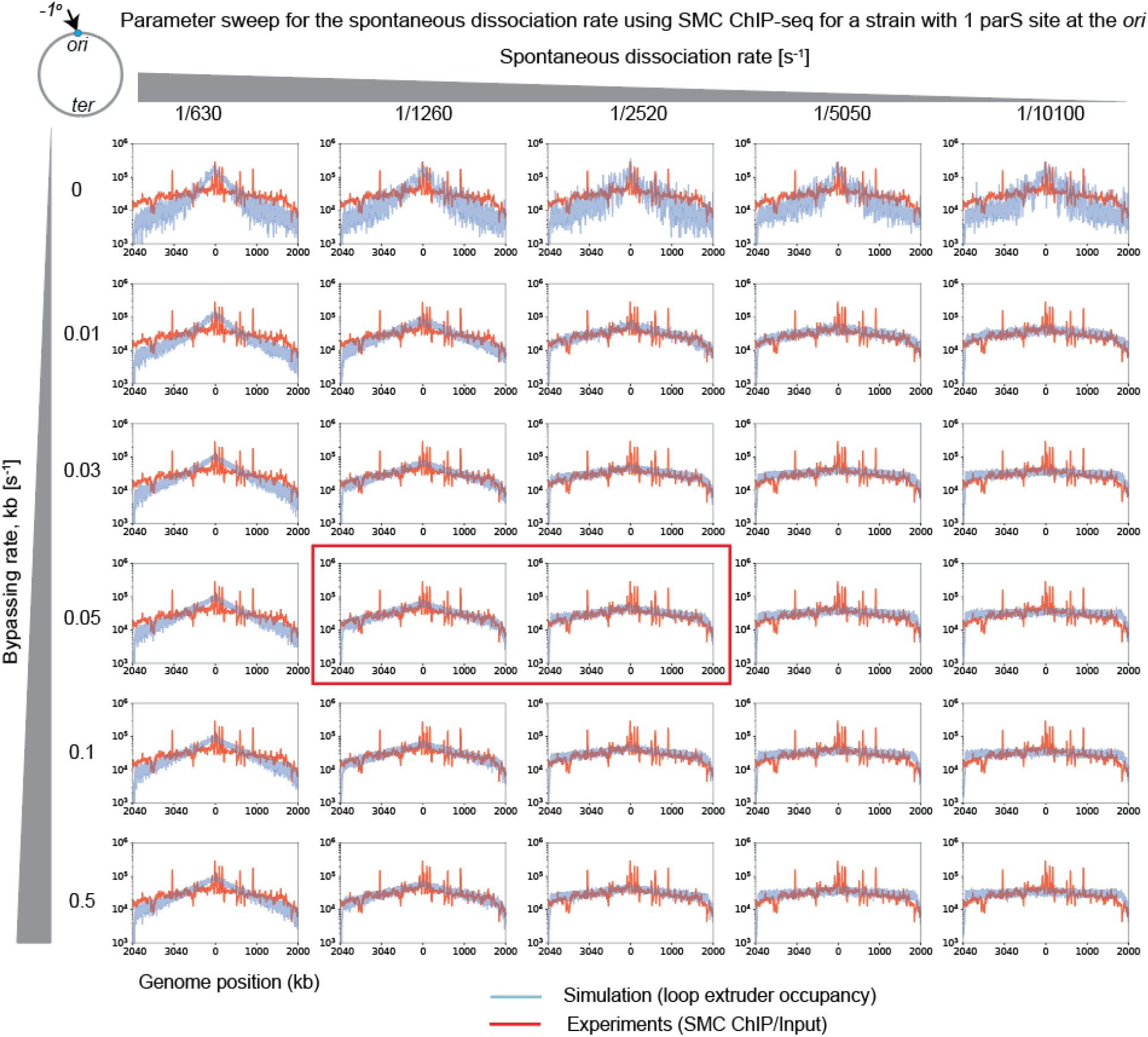
The spontaneous dissociation rate controls SMC complex abundance as a function of distance from the *parS* site. To obtain an estimate of the spontaneous dissociation rate, we compared the input-normalized experimental SMC ChIP-seq profile to the occupancy profile of loop extruders from simulation. We used a model where extrusion complexes can bypass one another and dissociate spontaneously, and where the facilitated unloading rate (from SMC complex encounters was set to zero). The experimental data was obtained from a strain with a single *parS* site near the origin (Wang et al, 2017), and was compared to a model with a *parS* site also at the −1° position. Notably, when the bypassing rate was 0 s^−1^, loop extruders accumulate strongly near the loading site for all values of the dissociation rate (i.e. top row). Additionally, if the dissociation rate was too high (≳ 1/630 s^−1^) or too low (≲ 1/5050 s^−1^), then the occupancy profile was too steep or too shallow, respectively. The optimal profiles (shown in the black box) were obtained for the bypassing rate near 0.05 s^−1^ and dissociation rates 1/1260 s^−1^ and 1/2560 s^−1^. We chose 1/2560 s^−1^ to be the default for all simulations thereafter as it also gave a better agreement with the Hi-C contact maps.

**Figure S9:**
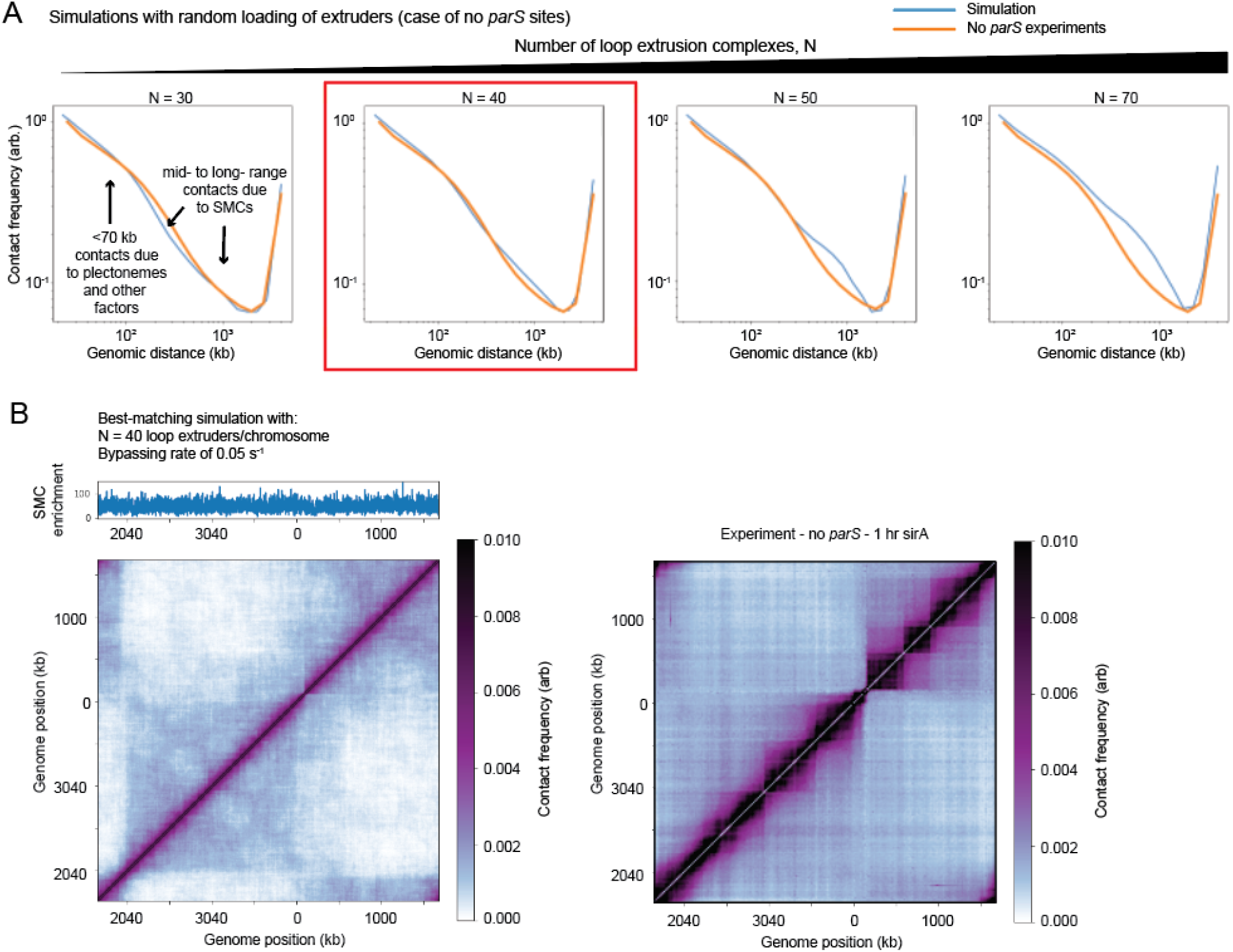
Determining the number of loop extruders per chromosome by matching contact probability decay curves. **(A)** Comparison of the contact probability decay curve of 3D polymer simulations with bypassing rates and varying numbers of extruders (as specified). The bypassing rate was constrained theoretically in relation to N, plectonemes were created with an average length of 45 kb and contacts were computed using a 9 monomer cutoff radius (i.e. ~ 9 kb) (see **Supplemental Information**). **(B)** Contact map and SMC enrichment profile for the simulation with N = 40 extrusion complexes (left) and experimental contact map (right).

**Figure S10:**
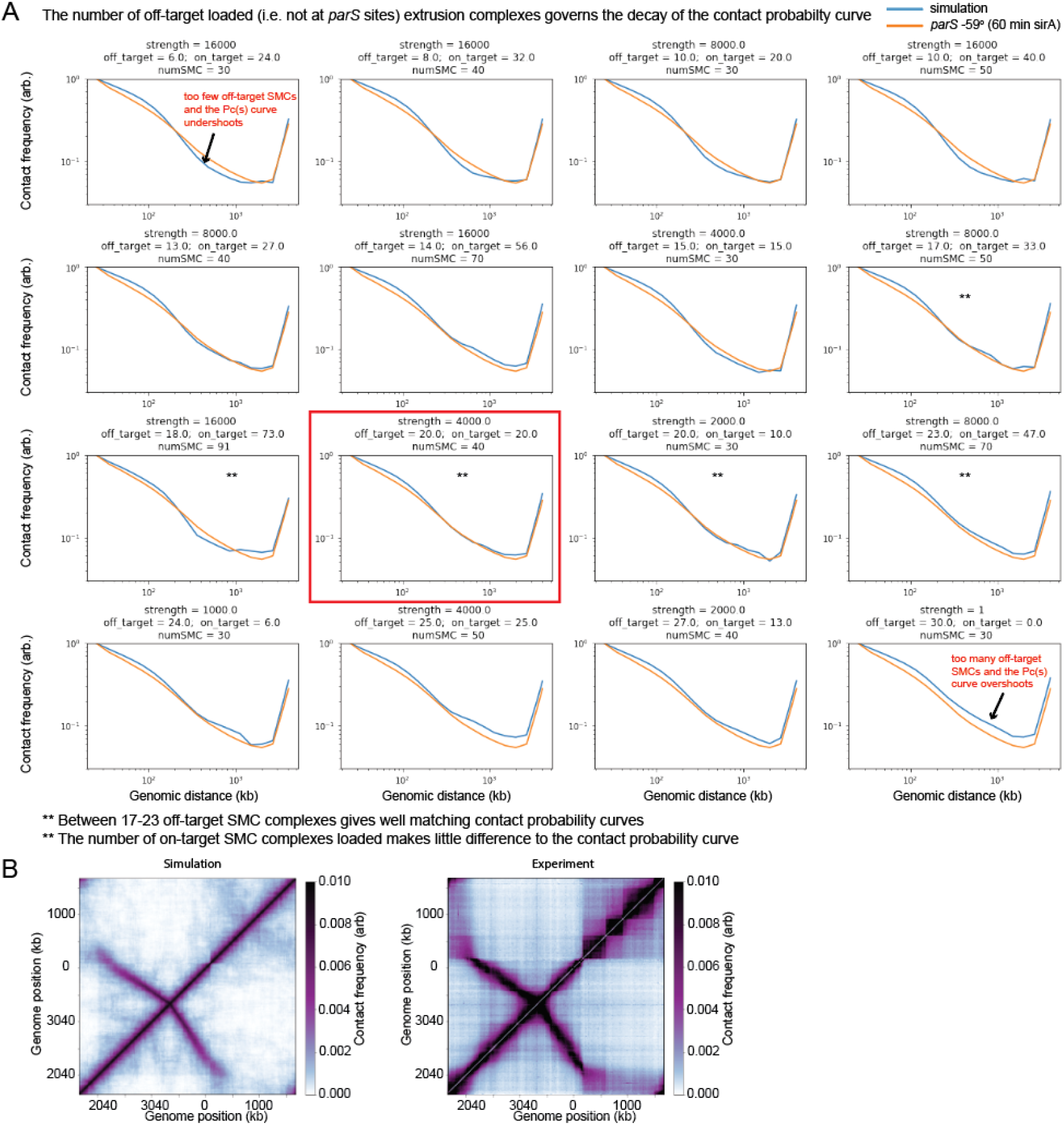
Determining the basal (non *parS* site to *parS* site) loading rate of SMCs using Hi-C from strains with a single *parS* site. Contact probability decay curves are shown for a parameter sweep of simulations where the rate of loading between *parS* and non-*parS* sites was varied. 3D polymer simulations were performed for a single *parS* site at the −59° position. Plectonemes of length 45 kb were included in these simulations to reproduce the experimental short-range contact probability. The bypassing rate was constrained theoretically in relation to N (see **Supplemental Information**). The strength of the *parS* site is shown above each graph denoted “strength”, indicating the relative likelihood that an SMC will load at the *parS* site monomer versus any other of the 4040 simulation monomers. The average number of extruders loaded at *parS* site versus off *parS* sites is indicated by “on target” and “off target”, respectively. The total number of extruders present in a simulation is indicated by the value “numSMC” = “on target” + “off target”. The black box highlights the best matching curves for simulations with a total number of extruders of N=40 (which was the best matching value for the strain with no *parS* sites, **see** **Fig. S9**). Thus, the best matched *parS* site monomer strength has a value of 4000-fold more than non-*parS* sites; with one *parS* site present on the genome, the SMC complexes load ~50% at the *parS* site and 50% elsewhere.

**Figure S11:**
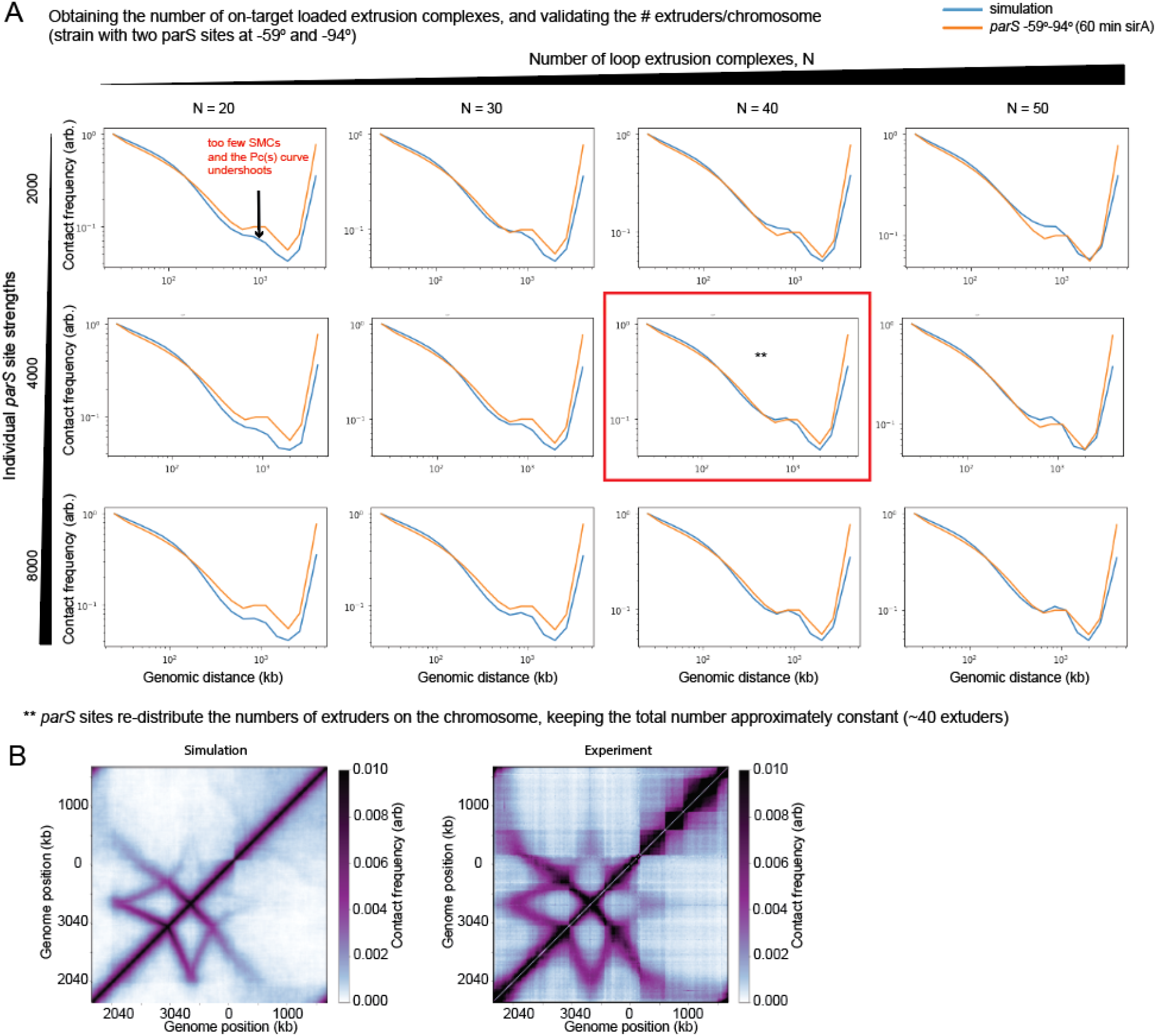
Verification and validation of the numbers of extruders present per chromosome and the *parS* loading rate. **(A)** Using the values identified for N in Fig. S9 and *parS* strength in Fig. S10, we obtain an excellent match to the contact probability decay curve for the strain harbouring two *parS* sites at −59° and −94°. We see that with two parS sites present on the genome, the SMC complexes load preferentially 66% at the *parS* sites and 33% elsewhere. **(B)** Comparison of the simulated and experimental Hi-C maps corresponding for the conditions and parameters shown in the red box in **A**.

**Figure S12:**
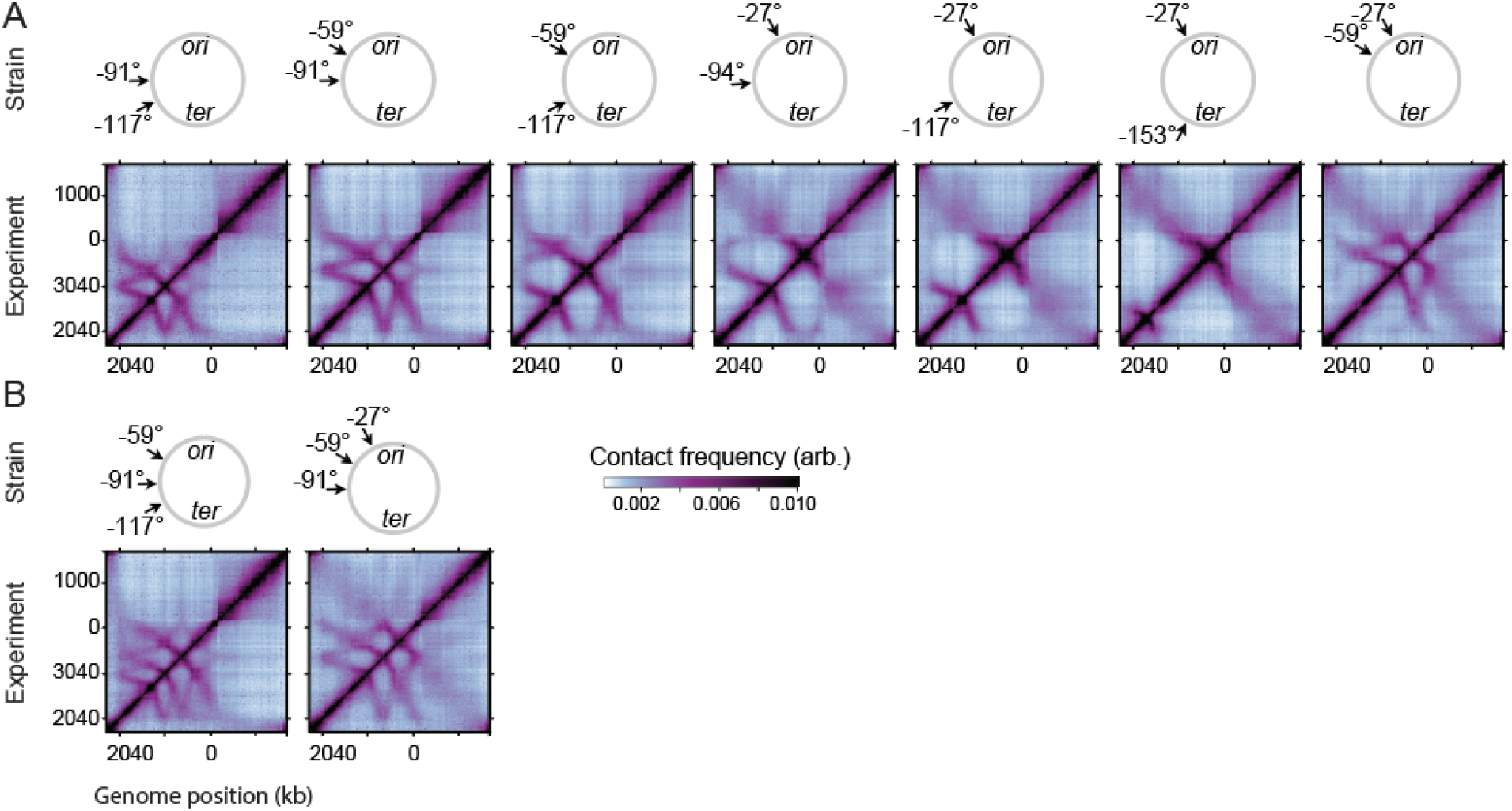
Hi-C maps for all stains in exponential growth. Hi-C maps for strains with **(A)** two *parS* sites, and **(B)** three *parS* sites.

**Figure S13:**
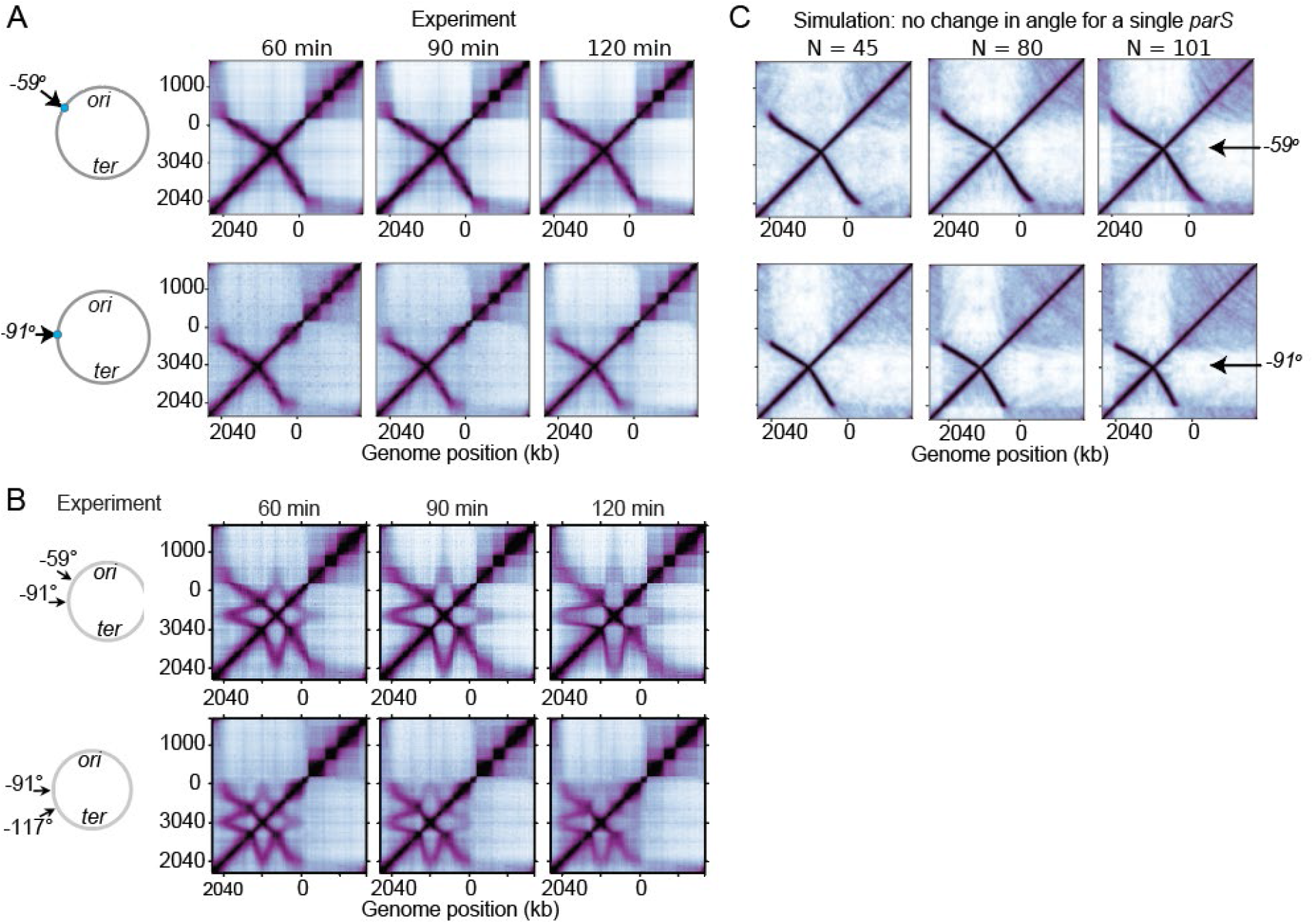
Experiment and model of the G1 arrest for strains with one or two *parS* sites. The experimental time course of G1 arrest for a strain with **(A)** a single *parS* site at the −59° position (top) and the −91° position (bottom), and (**B**) with two *parS* sites at −59° −91° (top) and −91° −117° (bottom). The experiments show that almost no change occurs to the angle of the hairpin structures when only a single *parS* site is present, but the hairpins increasingly tilt away from each other when two *parS* sites are present. **(C)** A 3D polymer simulation with the blocking, bypassing and unloading model of loop extrusion showing that when a single *parS* site is present, increasing the numbers of loop extruders, N, on the chromosome also does not change the observed hairpin angle for the same strains as in (A). Loop extrusion parameters use a bypassing rate of 0.05 s^−1^ and a facilitated dissociation rate of 0.003 s^−1^ (i.e. same as Fig. 4A); the number of extrusion complexes is denoted by N.

**Figure S14:**
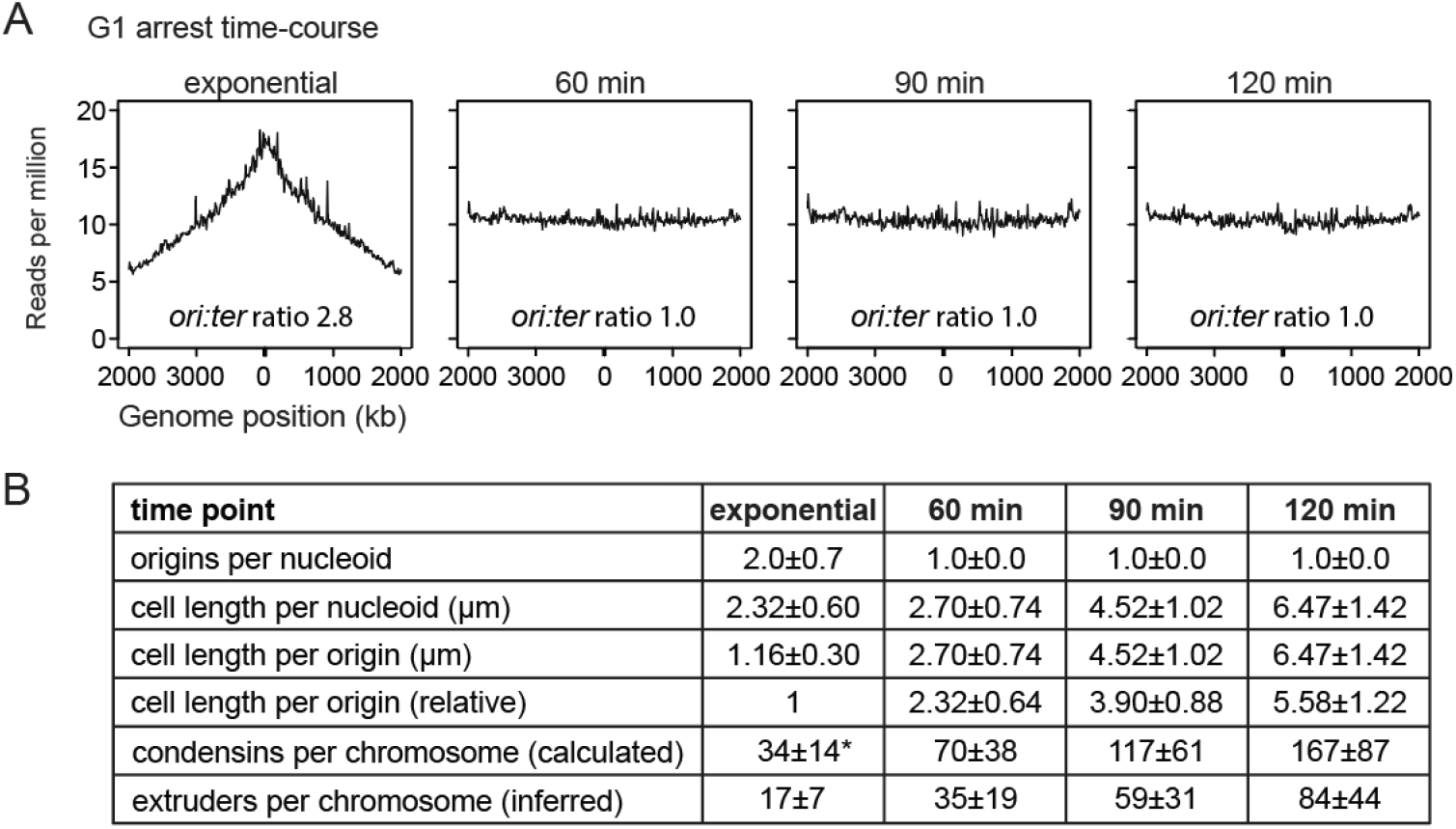
Quantification of chromosome numbers and cell lengths per nucleoid. **(A)** Whole genome sequencing for cells after the indicated minutes of replication inhibition by SirA. The computed *ori:ter* ratio indicates that by 60 min of SirA treatment, cells have finished chromosome replication. **(B)** The quantification of microscopy images reveals the numbers of origins per nucleoid, and cell lengths per nucleoid. These values are used to calculate the numbers of condensins per chromosome at different time points. To estimate the absolute numbers of condensins/chromosome (independently of the Hi-C data and polymer simulations), we use the reference value of 30 condensins/*ori* as measured in (Wilhelm et al, 2015), which converts to 34 condensins/chromosome (indicated by *). We infer that these calculated values agree well with the numbers of loop extrusion complexes (as found by Hi-C and polymer modeling), if there are two condensins per loop extrusion complex; this inference assumes that the error on the reference value of 30/condensins/*ori* is sufficiently small. For calculations, see the attached Excel file: Condensin numbers by MFA with error propagation (**Supplemental Information**).

**Figure S15:**
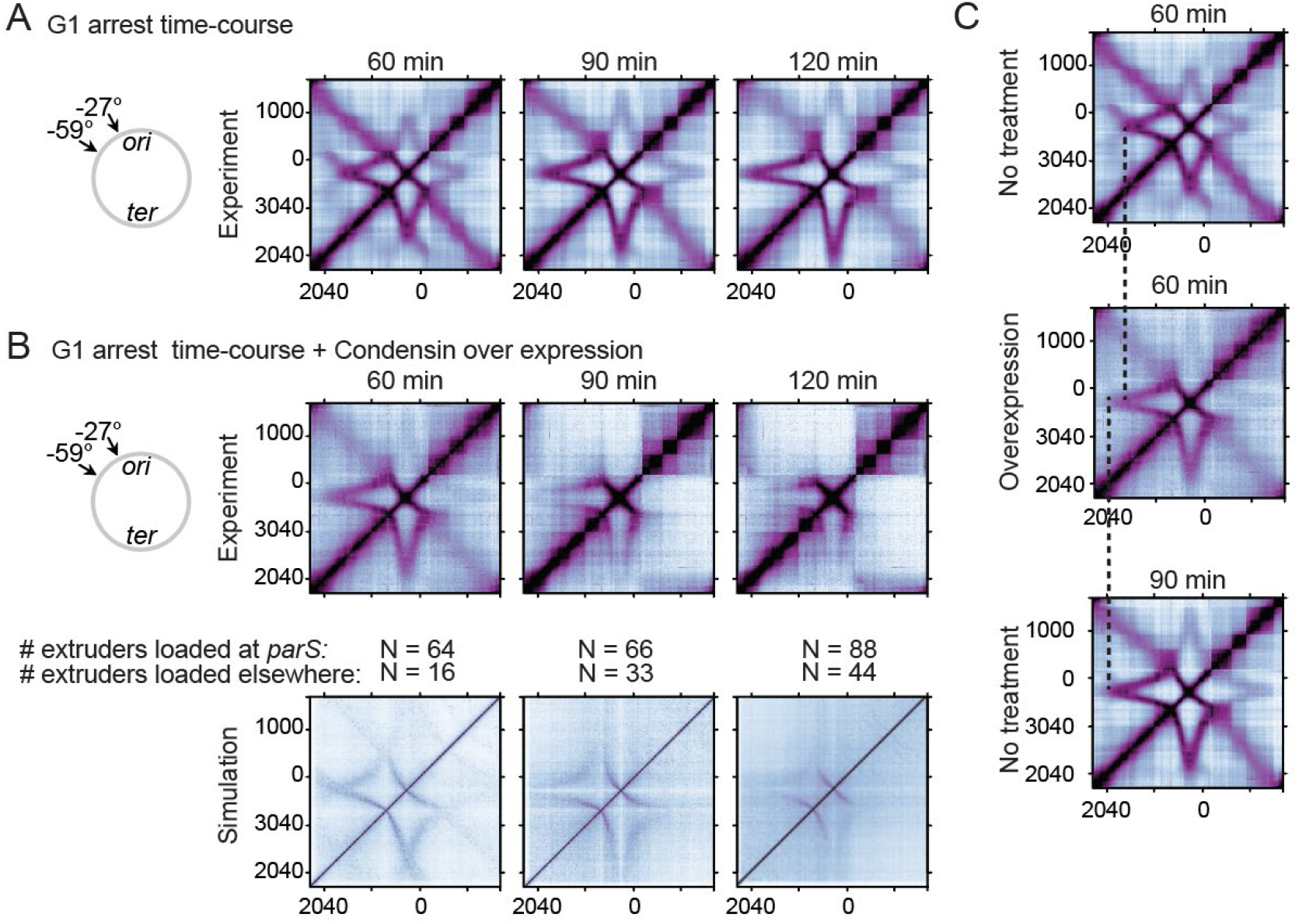
Overexpression of condensin speeds up effects of G1 arrest. **(A)** Replication inhibition Hi-C time course following induction of SirA for a strain with *parS* sites at −27° and −59°. (**B**) Condensin was overexpressed in the same background as the strain in panel A. We found that prolonged over-expression of SMC complexes at 90 min and 120 min did not recapitulate the experiments seen in G1 arrested cells (A) but caused the interaction lines to become shorter. These patterns are likely due to non-specific loading of SMC complexes outside of *parS*, creating traffic jams along the DNA. By simulations, we see that by increasing the numbers of off-*parS* loaded extruders, while keeping the numbers of on-*parS* loaded extruders consistent, we can observe similar changes in the Hi-C maps. Numbers of on-*parS* versus off-*parS* loading are average values for the simulation. (**C**) The size of the star-shaped pattern for the condensin overexpression strain at 60 min following SirA induction more closely resembles the pattern at 90 min with no overexpression (i.e. no treatment) than the corresponding 60 minute time-point with no overexpression. This indicates that increasing the numbers of condensins on the chromosome leads to an increase in the tilts of the hairpin diagonals away from each other.

**Figure S16:**
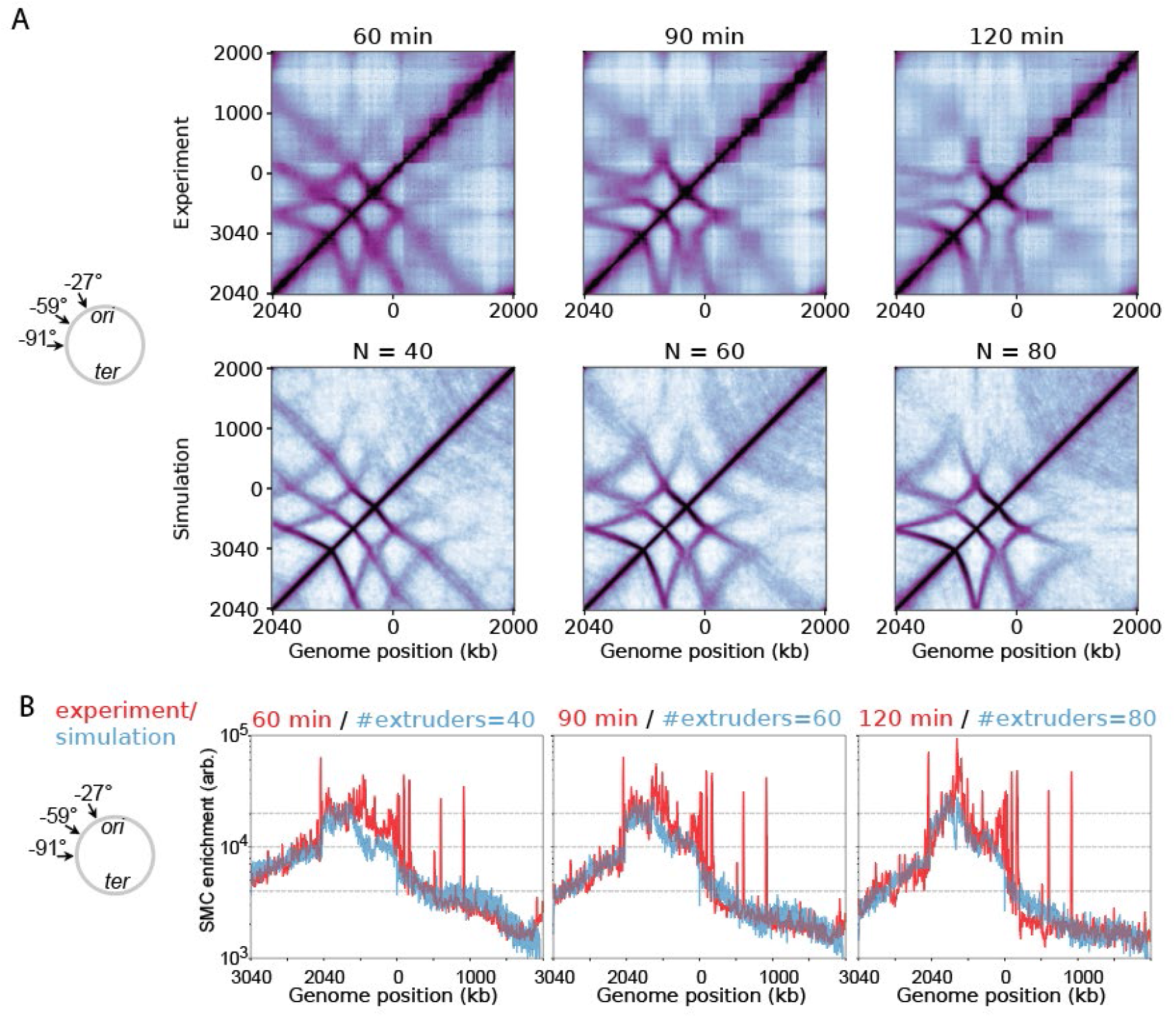
Experiment and model of the G1 arrest for strains with 3 *parS* sites. (**A**) Time-course Hi-C of SirA replication inhibition (and corresponding 3D polymer simulations) for a strain with three *parS* sites at positions −91°, −59°, −27°. (**B**) ChIP-seq performed for the same strains; experiments (red) and simulations (blue). (**A-B**) In the simulations, the bypassing rate was 0.05 s^−1^ and the facilitated dissociation rate was 0.003 s^−1^ (i.e. same as Fig. 4A).

**Figure S17:**
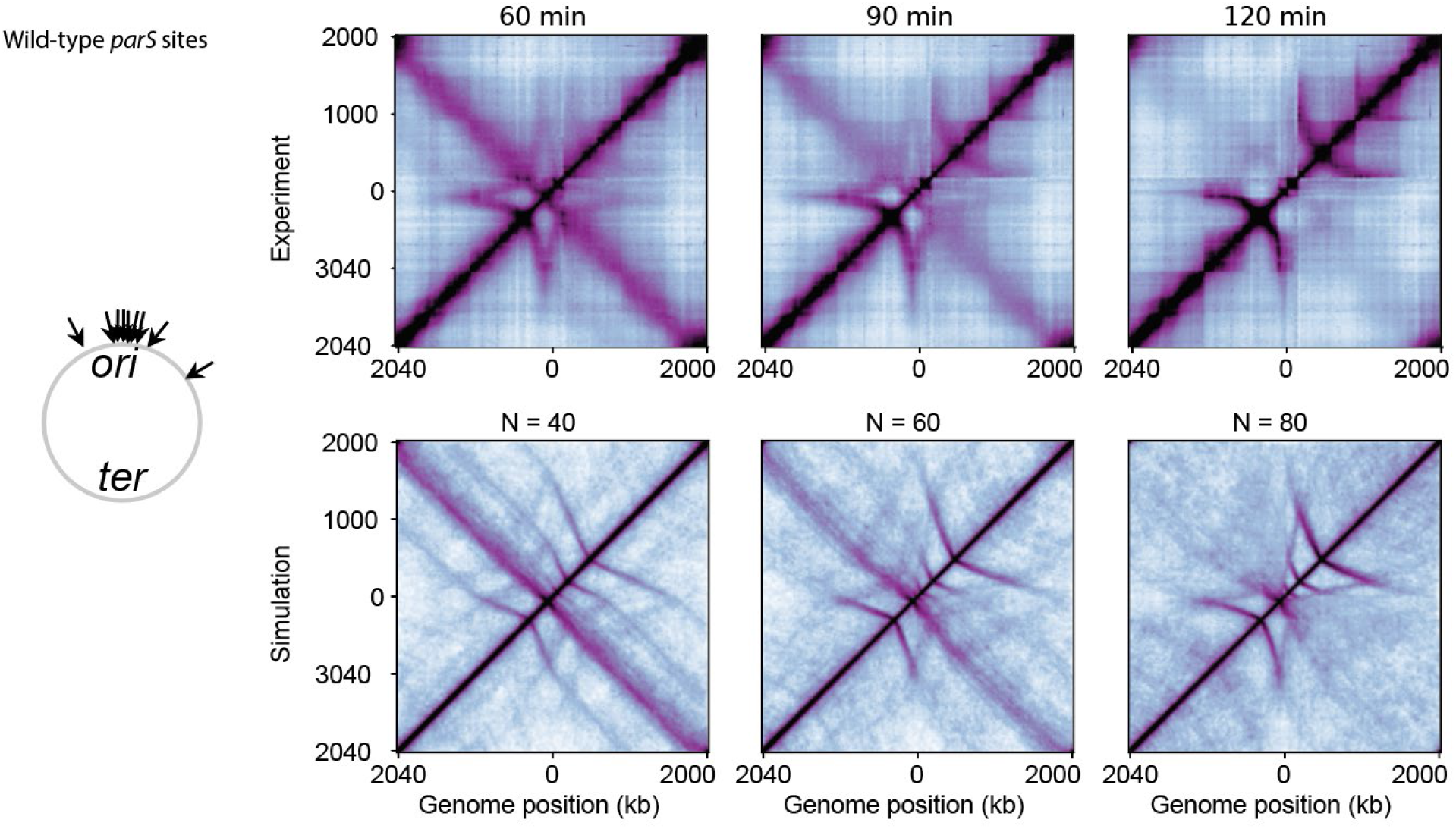
Experiment and model of the G1 arrest for strains with wild-type *parS* sites. The wild-type *parS* sites occur at the positions (−27°, −6°, −5°, −4°, −1°, +4°, +17°, +42°, +91°) in the strain *PY79*. In the simulation, we included −27°, −6°, −5°, −4°, −1°, +4°, +17°, +42°, +91° as extrusion from the position +91° is partially attenuated by a proximal chromosome interaction domain boundary and because the +91° *parS* site is weaker than the others (i.e. there is less ParB binding there). As in the experiments (top), in simulations (bottom) the central diagonal gradually vanishes due to traffic jams between extruders at the *ori-*proximal *parS* sites after increasing numbers of loop extruders. The bypassing rate was 0.05 s^−1^ and the unloading rate was 0.003 s^−1^ in the simulations.

**Figure S18:**
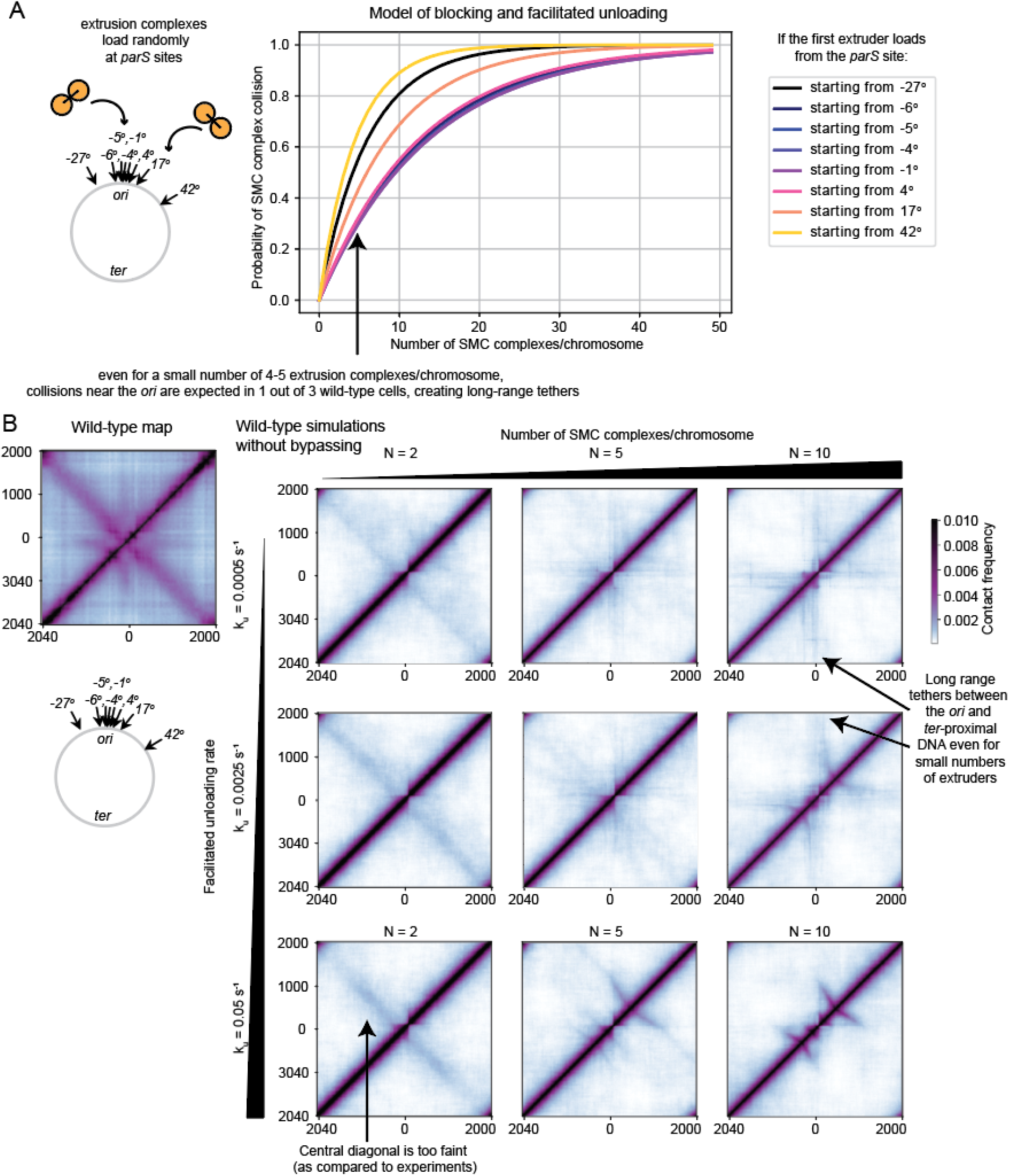
Blocking and unloading model applied to WT cells. **(A)** Analytical results demonstrating there is a high likelihood of collisions between SMC complexes near the *ori* due to the high density of *parS* sites. Calculations were performed for a facilitated unloading rate of 0.0006 s^−1^ and an extrusion rate of 0.8 kb/s. **(B)** 3D polymer simulations showing that even a few loop extruders (e.g. 5 extruders) results in a missing central diagonal and long-range tethers between the *ori* and other genome positions. With more extruders per chromosome, the traffic jams between SMC complexes near the origin becomes more likely, preventing juxtaposition of the arms. For very low numbers of extruders (e.g. 2 extruders), the central diagonal is present, but it is much fainter than observed experimentally.

## Materials and Methods

### General methods

*Bacillus subtilis* strains were derived from the prototrophic strain PY79 ^1^. Cells were grown in defined rich medium (CH) ^2^ at 37°C with aeration. Cells were arrested at the G1 phase by expressing SirA ^3^ for indicated durations using IPTG (isopropyl β-d-1-thiogalactopyranoside) at a final concentration of 1 mM or xylose at 0.5%. A list of Next-Generation Sequencing samples can be found in Table S1 arranged by the figure in which they appear. Lists of strains, plasmids and oligonucleotides can be found in Tables S2-S4.

### Hi-C

The detailed Hi-C procedure was previously described ^4^. Briefly, 5×10^7^ cells were crosslinked with 3% formaldehyde at room temperature for 30 min then quenched with 125 mM glycine. Cells were lysed using Ready-Lyse Lysozyme (Epicentre, R1802M) followed by 0.5% SDS treatment. Solubilized chromatin was digested with HindIII for 2 hrs at 37°C. The cleaved ends were filled in with Klenow and Biotin-14-dATP, dGTP, dCTP, dTTP. The products were ligated in dilute reactions with T4 DNA ligase overnight at 16°C. Crosslinks were reversed at 65°C overnight in the presence proteinase K. The DNA was then extracted twice with phenol/chloroform/isoamylalcohol (25:24:1) (PCI), precipitated with ethanol, and resuspended in 20 μl of QIAGEN EB buffer. Biotin from non-ligated ends was removed using T4 polymerase (4 hrs at 20°C) followed by extraction with PCI. The DNA was then sheared by sonication for 12 min with 20% amplitude using a Qsonica Q800R2 water bath sonicator. The sheared DNA was used for library preparation with the NEBNext UltraII kit (E7645) according to the manufacturer’s instructions for end repair, adapter ligation, and size selection. Biotinylated DNA fragments were purified using 10 μl streptavidin beads. 5 μl DNA-bound beads were used for PCR in a 50 μl reaction for 14 cycles. PCR products were purified using Ampure beads and sequenced at the Indiana University Center for Genomics and Bioinformatics using NextSeq 550. Paired-end sequencing reads were mapped to the genome of *B. subtilis* PY79 (NCBI Reference Sequence NC_022898.1) using the same pipeline described in Wang et al., 2015 ^4^. The *B. subtilis* PY79 genome was first divided into 404 10-kb bins. Subsequent analysis and visualization was done using R scripts. The genetic loci marked by degree (°) were calculated using the PY79 genome, which results in a slight shift from data published using *B. subtilis* 168 genomic coordinates.

### ChIP-seq

Chromatin immunoprecipitation (ChIP) was performed as described previously ^4^. Briefly, cells were crosslinked using 3% formaldehyde for 30 min at room temperature and then quenched, washed, and lysed. Chromosomal DNA was sheared to an average size of 250 bp by sonication using a Qsonica Q800R2 water bath sonicator. The lysate was then incubated overnight at 4°C with anti-SMC ^5^ antibodies, and was subsequently incubated with Protein A-Sepharose (GE HealthCare) for 1h at 4°C. After washes and elution, the immunoprecipitate was incubated at 65°C overnight to reverse the crosslinks. The DNA was further treated with RNaseA, Proteinase K, extracted with PCI, resuspended in 50 μl EB and used for library preparation with the NEBNext UltraII kit (E7645) and sequenced using the Illumina MiSeq or NextSeq550 platforms. The sequencing reads were aligned to the *B. subtilis* PY79 genome (NCBI NC_022898.1) using CLC Genomics Workbench (CLC Bio, QIAGEN), and subsequently normalized, plotted and analyzed using R scripts.

### Whole Genome Sequencing for DNA replication profiling

Cells were grown and collected at the indicated time points. Genomic DNA was extracted using the QIAgen DNeasy Blood and Tissue kit (QIAgen 69504). DNA was sonicated using Qsonica Q800R2 sonicator for 12 min at 20% amplitude, to achieve an average fragment size of 250 bp. DNA library was prepared using NEBNext UltraII kit (NEB E7645), and sequenced using Illumina NextSeq550. Sequencing reads were mapped to *B. subtilis* PY79 genome (NCBI Reference Sequence NC_022898.1) using CLC Genomics Workbench (QIAgen). The mapped reads were normalized to the total number of reads for that sample and plotted in R.

### Microscopy

Fluorescence microscopy was performed on a Nikon Ti2E microscope equipped with Plan Apo 100x/1.4NA phase contrast oil objective and an sCMOS camera. Cells were immobilized using 2% agarose pads containing growth media. Membranes were stained with FM4-64 (Molecular Probes) at 3 μg/ ml. DNA was stained with DAPI at 2 μg/ml. Images were cropped, adjusted and analyzed using MetaMorph software (Molecular Devices). Final figures were prepared in Adobe Illustrator.

### Immunoblot analysis

Cells were collected at appropriate time points and resuspended in lysis buffer (20 mM Tris pH 7.0, 1 mM EDTA, 10 mM MgCl_2_, 1 mg/ml lysozyme, 10 μg/ml DNase I, 100 μg/ml RNase A, 1 mM PMSF and 1% proteinase inhibitor cocktail (Sigma P-8340) to a final OD_600_ of 10 for equivalent loading. The cell resuspensions were incubated at 37°C for 10 min for lysozyme treatment, and followed by the addition of an equal volume of 2x Laemmli Sample Buffer (Bio-Rad 1610737) containing 10% β-Mercaptoethanol. Samples were heated for 5 min at 80°C prior to loading. Proteins were separated by precast 4-20% polyacrylamide gradient gels (Bio-Rad 4561096), electroblotted onto mini PVDF membranes using Bio-Rad Transblot Turbo system and reagents (Bio-Rad 1704156). The membranes were blocked in 5% nonfat milk in phosphate-buffered saline (PBS) with 0.5% Tween-20, and then probed with anti-ParB (1:5,000) ^6^, anti-SMC (1:5,000) ^5^, anti-SigA (1:10,000) ^7^, anti-ScpA (1:10,000) ^8^, or anti-ScpB (1:10,000) ^8^ diluted into 3% BSA in 1x PBS with 0.05% Tween 20. Primary antibodies were detected using Immun-Star horseradish peroxidase-conjugated goat anti-rabbit antibodies (Bio-Rad 1705046) and Western Lightning Plus ECL chemiluminescence reagents (Perkin Elmer NEL1034001) as described by the manufacturer. The signal was captured using ProteinSimple Fluorchem R system. The intensity of the bands were quantified using ProteinSimple AlphaView software.

### Plasmid construction

#### pWX512

[*amyE::Phyperspank-(optRBS)-smc (spec)*was generated by inserting *smc* with an optimal ribosome binding site (*optRBS*) (amplified using oWX516 and oWX517 from *B. subtilis* PY79 genome and digested with NheI and SphI) into pdr111 [*amyE::Phyperspan (spec)*] (D. Z. Rudner, unpublished) between NheI and SphI. The construct was sequenced using oWX486, oWX524, oWX848, oWX1194, oWX1195 and oWX1196.

#### pWX777

[*yhdG::Pxyl-(optRBS)-sirA (phleo)*] was generated by inserting *sirA* with an optimal ribosome binding site (*optRBS*) (amplified using oWX1892 and oWX1893 from *B. subtilis* PY79 genome and digested with HindIII and NheI) into pMS25 [*yhdG::Pxyl (phleo)*] (D. Z. Rudner, unpublished) between HindIII and NheI. The construct was sequenced using oML87 and oWX1894.

#### pWX778

[*yhdG::Phyperspank-(optRBS)-scpAB (phleo)*] was generated by inserting *scpAB* with an optimal ribosome binding site (*optRBS*) (amplified using oWX1897 and oWX1898 from *B. subtilis* PY79 genome and digested with HindIII and NheI) into pMS28 [*yhdG::Phyperspank (phleo)*] (D. Z. Rudner, unpublished) between HindIII and NheI. The construct was sequenced using oWX428, oWX486 and oWX487.

#### pWX788

[*yhdG::Phyperspank-(optRBS)-sirA (erm)*] was generated by inserting *sirA* with an optimal ribosome binding site (*optRBS*) (amplified using oWX1892 and oWX1893 from *B. subtilis* PY79 genome and digested with HindIII and NheI) into pMS24 [*yhdG::Phyperspank (erm)*] (D. Z. Rudner, unpublished) between HindIII and NheI. The construct was sequenced using oWX486 and oWX524.

### Strain construction

#### −91° *parS loxP-kan-loxP* (BWX3379)

The +4° *parS* sequence (TGTTACACGTGAAACA) was inserted at −91° (in the intergenic region between *ktrB* and *yubF*). An isothermal assembly product was directly transformed to *parSΔ9* (BWX3212) ^4^, which has all the 9 *parS* sites deleted from the *B. subtilis* genome. The isothermal assembly reaction contained 3 PCR products: 1) a region containing *ktrB* (amplified from PY79 genomic DNA using oWX1279 and oWX1280); 2) *loxP-kan-loxP* cassette flanked by the +4° *parS* sequence (amplified from pWX470 using universal primers oWX1241 and oWX438) and 3) a region containing *yubF* (amplified from PY79 genomic DNA using primers oWX1281 and oWX1282). The transformants were amplified and sequenced using oWX1283 and oWX1284.

Multiple *parS* sites were combined by standard transformation protocols. *loxP-kan-loxP* cassette was removed using a *cre* expressing plasmid pDR244 ^9^, resulting in an unmarked *parS* site indicated as “*no a.b.”.*

### Loop Extrusion Simulations

Numpy 1.18.1 was used for most of the calculations below ^10^.

#### Lattice set-up

We define the chromosome as a lattice of *L* = 4040 sites, where each lattice site corresponds to ~1 kb of DNA. The origin of replication is situated at lattice position 0, and the terminus is between lattice positions 1950-2050. Since the genome is circular, the lattice position of monomer 4040 is connected to the first monomer; as such, loop extrusion steps occur with periodic boundary conditions (i.e. a step from lattice site 4040 to 4041 becomes effectively a step from 4040 to 1).

#### Time steps and rates of extrusion

We use a fixed-time-step Monte Carlo algorithm as in previous work ^11^. The 1D extrusion simulations proceed with time-steps equivalent to 1/20^th^ of a second. Loop extruding factors (LEFs) are represented as two motor subunits, which move independently from one another in opposing directions one lattice site at a time ^12^. To account for the asymmetric rates of extrusion observed in *B. subtilis* experimentally ^8^, we introduce direction specific rates: Accordingly, a LEF subunit moving in the *ori* to *ter* direction will change lattice sites with a probability 0.05. This ensures that the LEFs have an average rate of extrusion (in the absence of interactions) of ~1 kb/s from *ori* to *ter*; in contrast, *a* LEF subunit moving in the *ter* to *ori* direction will take a step with a probability of 0.035. We define the “terminus” as being at position 2000 (i.e. in the middle of the 1950-2050 region).

#### Site-specific loading rates

We allow LEFs to randomly load at any lattice position. To mimic the effect of biased loading of LEFs at *parS* sites, we make loading at some lattice sites (those designated as *parS* sites) more probable. For example, to simulate a strain with a *parS* site at the −27-degree position on a real chromosome, we designate the lattice site position 3737 (i.e. (360-27)/360*4040 = 3737) to be the simulated *parS* location. We make the relative probability of loading at a *parS* lattice site as 4000 times stronger than non-*parS* sites. Thus, in a simulated strain with 1 *parS* site, the loading bias at the *parS* site is 4000 and ~50% of the LEFs will load at that position; for a strain with 2 *parS* sites, the total loading bias at *parS* sites is 8000 and 66% of LEFs will bind to *parS* site sites; for a strain with 3 *parS* sites, the total loading bias 12000 and ~75% of extruders load at *parS*, and so forth.

#### Site-specific and general dissociation rates

In addition to random loading, we simulate site-specific unloading. The terminus lattice sites (i.e. monomers 1950-2050) are given an unloading rate of 0.0025 s^−1^ (which is roughly ~5-fold higher than the basal dissociation rate). The basal dissociation rate is 0.0005 s^−1^. For each simulation time step, a random number is drawn between 0 and 1; if the value falls below the basal dissociation probability, then the LEF (i.e. both motor subunits) dissociate from the chromosome and load elsewhere.

#### Numbers of LEFs

We perform all simulations with fixed numbers of extruders. Thus, the effective loading and unloading rates on the chromosomes are always equal: when a LEF dissociates it associates immediately elsewhere on the genome.

#### Rules of LEF interaction (bypassing and unloading)

LEFs are deemed to encounter one another when they occupy adjacent lattice sites. By default, we do not allow LEF subunits to take a step onto an occupied site. The sole exception is if a LEF “bypasses” another; in this case, LEFs can move to co-occupy the same lattice site. Further steps may proceed unhindered if the adjacent sites are free.

To simulate bypassing and unloading we first define a bypassing probability, *b*, and an unloading probability, *u*, where b+u <= 1. Steps are taken following the principles of the Monte Carlo algorithm. At each simulation time step, a random number is drawn between 0 and 1. If the value is above (*b+u*), then the subunit location remains unchanged (i.e. no action is taken). If the value is below (*b+u*), but above *b,* then the LEF (i.e. both subunits) is marked to unload from the chromosome and re-load at the next time step following the loading rules above. Finally, if the value is below *b*, then the LEF subunit can move forward onto the occupied lattice site. Importantly, we note that at each simulation time step, *all* LEF subunits are updated simultaneously in this way. Moreover, the values *b* and *u*, will be slightly different depending on the direction of movement (i.e. *ori* to *ter*, or *ter* to *ori*, see above).

#### Loop extrusion equilibration steps

We compute 100,000 initialization steps for the loop extrusion simulations to ensure that the loop statistics have reached a steady-state before creating any contact maps.

### 3D Polymer Simulations

We coupled each of the 1D loop-extrusion simulations to a model of a polymer chain ^11^ and performed molecular dynamics simulations using Polychrom ^13^, a Python API that wraps the molecular dynamics simulation software OpenMM ^14^. In this coupled model, LEFs act as a bond between the two monomers. These bonds are dynamically updated depending on the position of the LEFs on the lattice (see “Loop Extrusion Simulations” section). From the polymer simulations, we obtain 3D polymer structures from which we can create contact frequency (Hi-C-like) maps (see below).

Polymers are constructed of *L* = 4040 consecutive monomers bonded via the pairwise potential:

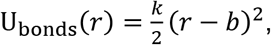

Where *k* = 2*k*_*b*_*T*/*δ*^2^ is the spring constant (*k*_*b*_ is the Boltzmann constant, *T* is the temperature, and *δ* = 0.1 is effectively the bond wiggle distance in monomer units), *r* = |*r*_*i*_ − *r*_*j*_| is the spatial displacement between monomers, and *b* = 1 is the mean distance between monomers in monomer units (typically ~30 nm). Monomers crosslinked by a LEF are held together by the same potential.

To account for excluded volume interactions between monomers, we have a weak polynomial repulsive potential:

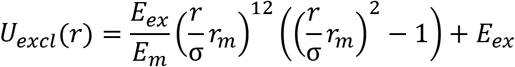

Defined for *r* < σ = 1.05, where 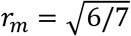, *E*_*m*_ = 46656/823543 and *E*_*ex*_ = 1.5*k*_*B*_*T*.

At the start of each simulation, the polymer is initialized as a random walk, with normally distributed velocities such that the total temperature is T. The system thermostat is set with an error tolerance of 0.01. Time steps integration is performed using the “variableLangevin” algorithm, and the collision rate is set to 0.03. Simulations were performed with periodic boundary conditions, where the box size was 27.2 monomer units in each dimension.

### Contact Map Generation from Simulations

Contact maps were obtained from loop extrusion simulations by two different methods: either using 3D polymer simulations, or by using a semi-analytical approach.

#### From 3D polymer simulations

The contact maps from 3D polymer simulation are displayed in both main text figures and supplemental figures; we used a distance cutoff radius of 9 monomers (or equivalently ~270 nm) and a minimum of 3,000 chromosome conformations to compute the contact maps. We note that the choice of cutoff radius did not significantly affect the observed contact patterns, but simply changed the perceived level of smoothness of the features. After an initial energy minimization and a further 6000 polymer simulation steps, we started recording chromosome conformations. Chromosome conformations were saved every 3000 polymer simulation steps, where each polymer simulation step contained 20 sub-steps of the 1D loop extrusion simulation.

#### From the semi-analytical approach

In addition to 3D polymer simulations, we generated contact maps semi-analytically. The semi-analytical approach employs a Gaussian approximation to calculate contact probability maps directly from the lattice positions of loop extruders. This approach allowed us to swiftly explore a broad range of model parameter values and generate Hi-C-like maps by circumventing the time-intensive 3D polymer simulations. We used a similar approach as previously ^11^ but with an extension to allow for z-loop like structures, which we explain below.

To compute the semi-analytical contact maps, we first create a non-directed graph of connections between monomers. Nodes of the graph represent monomers and edges represent connections between them. The graph contains edges between all nodes with indices (j, j+1); this creates the polymer chain backbone. Since the chromosome is circular, there is also an edge between the first and last node. Additional edges are introduced between all nodes (pairs of monomers) connected by a loop extruding factor. Thus, for a polymer chain of length L monomers with N extruders, the graph should contain L nodes, and L+N edges. The effective genomic separation between any two monomers (i,j) is obtained by computing the shortest path, *s*, between the monomers; we use Scipy’s ^15^ shortest_path function (SciPy 1.5.0) found in the scipy.sparse.csgraph module to find, *s*. The contact probability between these two points is then evaluated as s^(−3/2).

Contact probability maps were generated from at least 9,480,000 unique pairs of monomers. This represents 3,000 different chromosome conformations (i.e. different conformations of loop extruders), and 3160 unique samples from each conformation. For each chromosome conformation, we sampled contact probabilities by drawing a random list of 80 (out of 4040) monomers; we then computed the contact probability (as described above) between all unique monomer pairs (i.e. 80*79/2 = 3160 pairs) and stored this probability into a matrix. By repeating the process for each of the 3,000 chromosome conformations, and averaging the resulting probability matrices, we obtained a population averaged contact probability map.

For exact contact probability calculations, without the shortest path approximation, please see Banigan et al ^11^. We note, however, that the previously developed approach – while exact – does not account for z-loops (or pseudo-knots formed by extruders). It is thus not applicable to the simulations with bypassing extrusion. Moreover, while the shortest path approximation could affect the contact probabilities up to a factor of 2^(3/2)≈2.8 (i.e. the effective distance between the furthest points on a loop), it averages to an underestimation of contact frequency by a factor of ~1.5. As such, while we generally did not use the absolute values of the semi-analytically derived contact probabilities to draw quantitative conclusions about the Hi-C intensities, the semi-analytical maps could be used as an exploratory tool and to build intuition for the system.

#### Short-range contacts

To obtain a quantitative match between the contact probability decay curves from simulations and experiments, we needed to account for the shallow decay of contact probability at short distances (of lengths <60 kb). It was previously shown in *Caulobacter crescentus* that adding plectonemes of length ~30 kb was sufficient to get a match between polymer simulations and Hi-C data (Le et al, 2013). As proxy for supercoiling, we added a series of nested loop structures of average length 45 kb to our simulations all throughout the genome. They were constructed as follows: First, we generated a sorted list of 90 random integers between 1 and 4040 (corresponding to the lattice site positions). We added edges connecting the first and second, the third and fourth, the fifth and sixth, and so forth. This created a series of loops of average length 45 kb, separated by gaps of length 45 kb. We stored the positions of these additional “bonds” in a list. We repeated this process of generating a sorted list of 90 random integers from 1 to 4040 and creating edges. These two lists were appended together. This process produced overlapping loops of length 45 kb, mimicking the effect of supercoils. We generated new short-range contacts in this way for every 3000 polymer simulation steps, to randomize the short-range contact positions. These imposed contacts did not interfere with the lattice dynamics of the LEFs described above.

### SMC occupancy profiles from simulations

To compute the SMC occupancy profiles from the simulations, we captured at least 3,000 different LEF conformations. The temporal sampling of LEF conformations proceeded identically to the sampling of 3D polymer conformations (see “Contact map generation from simulations” above). To record the LEF occupancies, we created an array of length L=4040 bins (i.e. the same size as the chromosome) and added +1 counts to each bin occupied by each LEF subunit (note that each LEF has two subunits). Thus, if there were 40 LEFs present on the chromosome, then at each sample a total of +80 counts would be added to the array. To directly compare the LEF occupancy profiles to the normalized SMC ChIP-seq experimental data (i.e. ChIP/input), we computed the median ChIP/input value from the SMC ChIP-seq tracks and normalized the LEF occupancy to match the experimental median value.

### Condensin number calculations: error estimation

Refer to the attached Excel spreadsheet for the details on the condensin number calculations as derived by marker frequency analysis, fluorescence microscopy and quantitative immunoblotting.

### Theory

#### Relationship between the tilt of Line 1 (or Line 2) and the extrusion speeds between the *parS* sites

We derive in the sections below the relationship between the bypassing rates, numbers of extruders and the tilt of Line 1. However, the same procedure can be analogously applied to Line 2 arriving at similar answers.

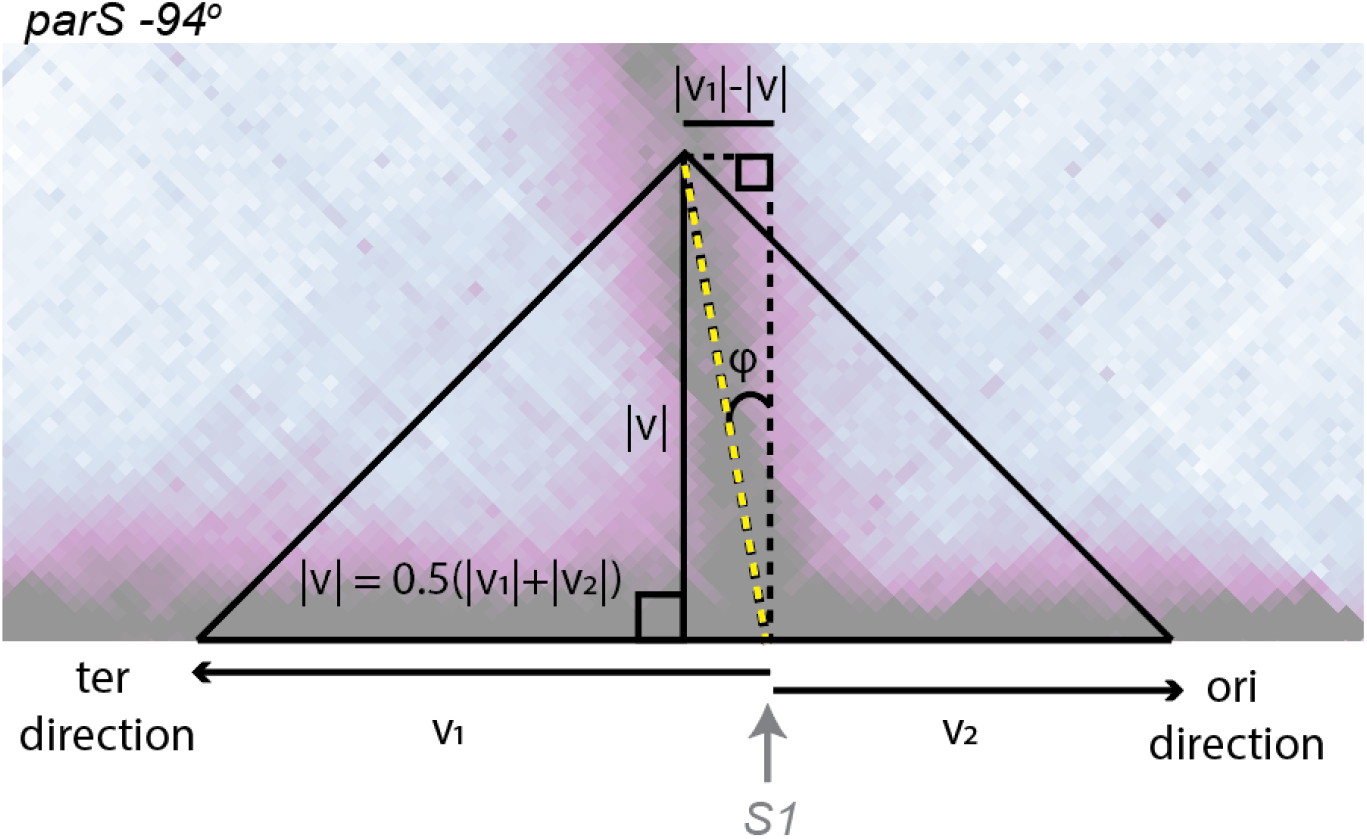

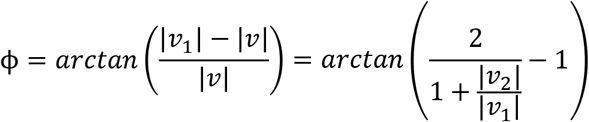

##### For a strain with one parS site at the −94° position (i.e. S1)

From the diagram and equation above (see also Fig. S2C) that the tilt of Line 1 is related to the relative LEF subunit translocation speeds towards *ori* versus *ter*. By our measurement of ϕ = 9.7^0^, we find that v_2_ ≈ 0.707 v_1_.

##### For a strain with two parS sites at the −94° and −59° position

Measuring the tilt of Line 1 in the strain with two *parS* sites (see below), we find and angle ϕ = 15.9^0^, or equivalently that v_3_ ≈ 0.556 v_1_.

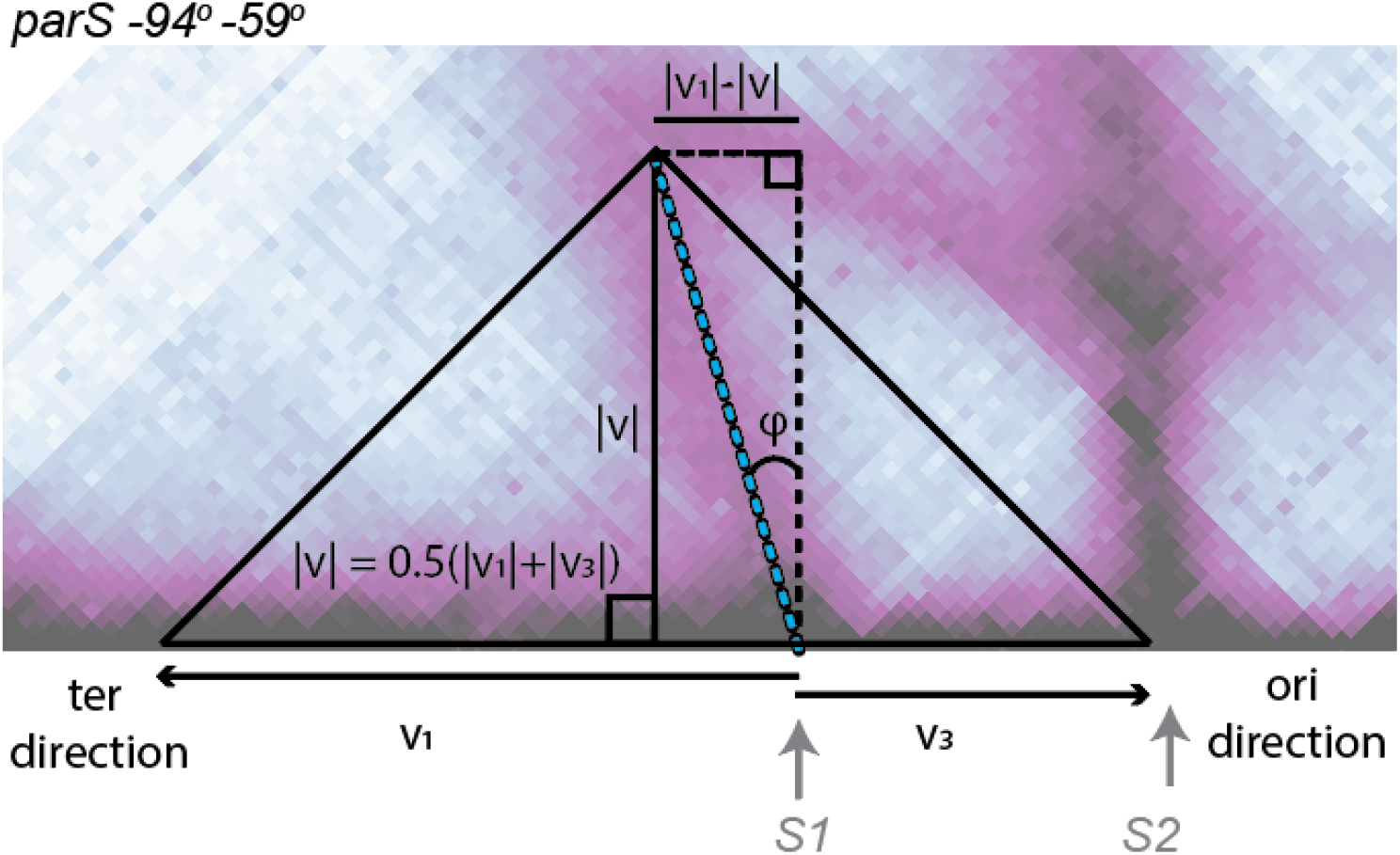

Hence, when there is more than 1 *parS* site, the average speed from *S1* to *S2* decreases by a factor of *v*_2_/*v*_3_ ≈ 1.2. Moreover, from the from the reference value *v*_1_ = 0.83 ± 0.17 kb/s from Fig. S2A and measurements in Wang et al, 2017 ^8^, we obtain that v_2_ = 0.59 ± 0.12 kb/s and *v*_3_ = 0.46 ± 0.09 kb/s.

#### Relationship between the bypassing rate, number of loop extruders and the tilt of Line 1

We can use the measured speeds v_2_ and v_3_ to estimate the average bypassing rate with a simple model: Consider a LEF subunit translocating a distance *d* from the *parS* site *S1* towards *S2*. We can define the time, τ, as the time it takes to move one lattice site of length *l*_0_ = 1 kb, if the adjacent lattice site unoccupied. We define τ_*b*_ as the time to bypass a lattice site occupied by another LEF. If, on average, the LEF subunit travelling from *S1* to *S2* encounters *n* other LEF subunits, then the total time, *T*_3_ to cross the distance *d* is:

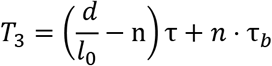

We can re-arrange the equation to obtain:

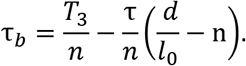

Noting that since *T*_3_ = *d*/*v*_3_ and τ = *l*_0_/*v*_2_, then:

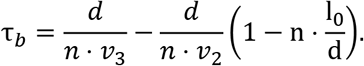

This helps constrain the parameter space for the search of best-fit parameters since the number of extruders and bypassing rate are linearly dependent on one another.

Moreover, we see that the product of the bypassing rate and the number of extruders per chromosome is approximately constant. For 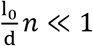, as reasonably expected for *n* < 100 (i.e. less than 200 loop extruders/chromosome), then:

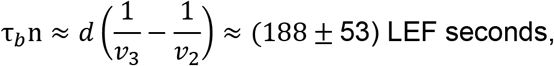

where we used known values: *d* = 392 kb for the distance between the two *parS* sites (in the −94° −59° strain) and the speeds v_2_ = 0.59 ± 0.12 kb/s and *v*_3_ = 0.46 ± 0.09 kb/s.

#### Estimating the bypassing rate independently of Hi-C and polymer simulations

We can calculate the bypassing rate, τ_*b*_, if we can estimate the number of extruders, n, that a LEF is expected to encounter on its transit from *S1* to *S2* using the relation τ_*b*_ ≈ 188/n seconds derived above.

For this calculation, we will need to know the number of LEFs moving from S2 to S1 when extrusion from S1 begins; additionally, we will need to know the number of LEFs that bind to S2 after extrusion from S1 has begun. Thus,

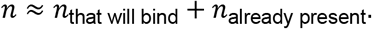

To calculate *n*_that will bind_, we will need to know the average time, *T*_3_, that it takes an extruder to cross the distance d (from S1 to S2) and the loading rate, *k*_*load*_, at the *S2 parS* site. It follows that:

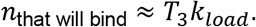

The value T_3_ = *d*/*v*_3_ ≈ 180 ± 35 *s*, where *d* is the distance between the two *parS* sites (i.e. 392 kb) and v_3_ is the average extrusion speed from *S1* to *S2* calculated above. We can calculate *k*_*load*_ from the total number of loop extruders per chromosome, *N*, and the dissociation rate, *k*_*d*_, of extruders using the relation:

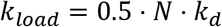

The number of extruders, *N*, is estimated to be *N* = (70±38)/*q*; extruders/chromosome where q=1 or 2 if extruders are monomers or dimers of condensins, respectively (see Fig. S14B). The factor of 0.5 in *k*_*load*_ arises if we assume that *S1* and *S2* have equal likelihood of loading the extruder.

The dissociation rate, *k*_*d*_, can be estimated from the average time it takes a LEF subunit to reach the terminus. As a first approximation, we can assume that a LEF immediately dissociates from the chromosome if any subunit reaches the terminus at genome position ~2000 kb; this is an acceptable assumption since experimentally SMCs do not accumulate at *ter* ^8^. After dissociating from *ter*, the LEF may re-load at either *S1* or *S2*. LEFs loaded at S1 travel a distance of ~980 kb to reach *ter* and take approximately (1180 ± 241) *s*. LEFs loaded at S2 travel a total distance of ~1372 kb to reach the terminus and take approximately (1845 ± 532) *s*. Thus, the average dissociation rate (per LEF), is then:

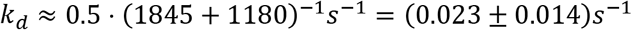

and

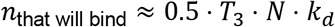

Combining all the above values and propagating the uncertainties, we obtain the estimate for the number of LEFs encountered as:

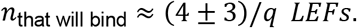

Next, we compute the number of LEFs moving from S2 to S1 that were already present in the segment between the *parS* sites at the time the extrusion from S1 began. We use the distance-weighted average:

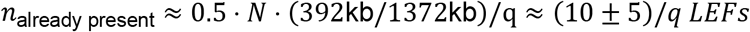

Finally, we obtain:

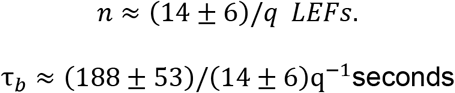

Thus, if loop extrusion complexes are made of condensin dimers, τ_*b*_ = 26 ± 19 *s*; if loop extrusion complexes are made of are condensin monomers, τ_*b*_ = 13 ± 9 *s*. These values are in good agreement with the average bypassing time obtained by 3D polymer simulations of τ_*b*_ = (20 ± 10) s, and measurements of the bypassing time obtained *in vitro* of ~8 seconds ^16^.

#### The frequency of nested doublet interactions is controlled by the ratio of bypassing to unloading rates

Nested doublet configurations (Fig. 2A) occur when a LEF from one *parS* site (e.g. S1) extrudes past the other site (e.g. S2) followed by a LEF loading event (i.e. at S2). The frequency with which LEFs will enter into this configuration will depend critically on the bypassing and unloading rates as LEFs encounter one another. If the bypassing rate is 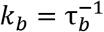 and the unloading rate is 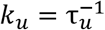 (i.e. due to collisions), then the probability that a LEF translocating from the S1 site manages to reach the S2 site (neglecting the basal dissociation rate) is given by:

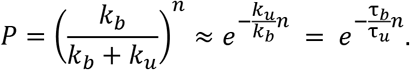

Where *n* is the expected number of encounters that a LEF translocating from S1 will have with other LEFs on its way to S2 (see calculation above).

From the relative intensity of Line 3 and Line 4 (from Hi-C data), we estimate that *P* ≈ 0.5. Using the values obtained above for *n*, it follows that τ_*u*_ ≈ 20τ_*b*_, meaning that the bypassing rate is 20-fold higher than the unloading rate.

**Supplemental Table S1.** Next-Generation Sequencing samples used in this study.

| figure | sample | data type |
| --- | --- | --- |
| Figure 1C (left) | 301_Wang_HiC_BWX4476_1mM_060m | Hi-C |
| Figure 1C (center) | 302_Wang_HiC_BWX4475_1mM_060m | Hi-C |
| Figure 1C (right) | 303_Wang_HiC_BWX4463_1mM_060m | Hi-C |
| Figure 1D (top, 0 min) | 304_Wang_HiC_BWX4493_xyl30m_1mM_00m | Hi-C |
| Figure 1D (top, 10 min) | 305_Wang_HiC_BWX4493_xyl30m_1mM_10m | Hi-C |
| Figure 1D (top, 15 min) | 306_Wang_HiC_BWX4493_xyl30m_1mM_15m | Hi-C |
| Figure 1D (top, 20 min) | 307_Wang_HiC_BWX4493_xyl30m_1mM_20m | Hi-C |
| Figure 1D (top, 25 min) | 308_Wang_HiC_BWX4493_xyl30m_1mM_25m | Hi-C |
| Figure 1D (top, 30 min) | 309_Wang_HiC_BWX4493_xyl30m_1mM_30m | Hi-C |
| Figure 1D (middle, 0 min) | 310_Wang_HiC_BWX4491_xyl30m_1mM_00m | Hi-C |
| Figure 1D (middle, 10 min) | 311_Wang_HiC_BWX4491_xyl30m_1mM_10m | Hi-C |
| Figure 1D (middle, 15 min) | 312_Wang_HiC_BWX4491_xyl30m_1mM_15m | Hi-C |
| Figure 1D (middle, 20 min) | 313_Wang_HiC_BWX4491_xyl30m_1mM_20m | Hi-C |
| Figure 1D (middle, 25 min) | 314_Wang_HiC_BWX4491_xyl30m_1mM_25m | Hi-C |
| Figure 1D (middle, 30 min) | 315_Wang_HiC_BWX4491_xyl30m_1mM_30m | Hi-C |
| Figure 1D (bottom, 0 min) | 316_Wang_HiC_BWX4492_xyl30m_1mM_00m | Hi-C |
| Figure 1D (bottom, 10 min) | 317_Wang_HiC_BWX4492_xyl30m_1mM_10m | Hi-C |
| Figure 1D (bottom, 15 min) | 318_Wang_HiC_BWX4492_xyl30m_1mM_15m | Hi-C |
| Figure 1D (bottom, 20 min) | 319_Wang_HiC_BWX4492_xyl30m_1mM_20m | Hi-C |
| Figure 1D (bottom, 25 min) | 320_Wang_HiC_BWX4492_xyl30m_1mM_25m | Hi-C |
| Figure 1D (bottom, 30 min) | 321_Wang_HiC_BWX4492_xyl30m_1mM_30m | Hi-C |
| Figure 2A, C | 303_Wang_HiC_BWX4463_1mM_060m | Hi-C |
| Figure 3A (column 1) | 322_Wang_HiC_BWX4462_1mM_060m | Hi-C |
| Figure 3A (column 2) | 323_Wang_HiC_BWX4479_1mM_060m | Hi-C |
| Figure 3A (column 3) | 324_Wang_HiC_BWX4480_1mM_060m | Hi-C |
| Figure 3A (column 4) | 325_Wang_HiC_BWX4481_1mM_060m | Hi-C |
| Figure 3A (column 5) | 326_Wang_HiC_BWX4883_1mM_060m | Hi-C |
| Figure 3A (column 6) | 327_Wang_HiC_BWX4482_1mM_060m | Hi-C |
| Figure 3A (column 7) | 328_Wang_HiC_BWX4892_1mM_060m | Hi-C |
| Figure 3B (row 1) | 329_Wang_HiC_BWX4927_1mM_060m | Hi-C |
| Figure 3B (row 2) | 330_Wang_HiC_BWX5066_1mM_060m | Hi-C |
| Figure 3C | 333_Wang_ChIPSMC_BWX4462_1mM_060m | ChIP-seq |
| Figure 3C | 336_Wang_input_BWX4462_1mM_060m | WGS |
| Figure 4A (column 1) | 322_Wang_HiC_BWX4462_1mM_060m | Hi-C |
| Figure 4A (column 2) | 331_Wang_HiC_BWX4462_1mM_090m | Hi-C |
| Figure 4A (column 3) | 332_Wang_HiC_BWX4462_1mM_120m | Hi-C |
| Figure 4B (row 1) | 333_Wang_ChIPSMC_BWX4462_1mM_060m | ChIP-seq |
| Figure 4B (row 2) | 334_Wang_ChIPSMC_BWX4462_1mM_090m | ChIP-seq |
| Figure 4B (row 3) | 335_Wang_ChIPSMC_BWX4462_1mM_120m | ChIP-seq |
| Figure 4B (row 1) | 336_Wang_input_BWX4462_1mM_060m | WGS |
| Figure 4B (row 2) | 337_Wang_input_BWX4462_1mM_090m | WGS |
| Figure 4B (row 3) | 338_Wang_input_BWX4462_1mM_120m | WGS |
| Figure S1B (left) | 29_Rudnerlab_HindIII_HiC_BWX3221 (GSE68418) <sup>4</sup> | Hi-C |
| Figure S1B (middle) | 08_Wang_HiC_BWX3377 (GSE85612) <sup>8</sup> | Hi-C |
| Figure S1B (right) | 339_Wang_HiC_BWX4428 | Hi-C |
| Figure S2A (row 1, column 1) | 310_Wang_HiC_BWX4491_xyl30m_1mM_00m | Hi-C |
| Figure S2A (row 1, column 2) | 311_Wang_HiC_BWX4491_xyl30m_1mM_10m | Hi-C |
| Figure S2A (row 1, column 3) | 312_Wang_HiC_BWX4491_xyl30m_1mM_15m | Hi-C |
| Figure S2A (row 1, column 4) | 313_Wang_HiC_BWX4491_xyl30m_1mM_20m | Hi-C |
| Figure S2A (row 1, column 5) | 314_Wang_HiC_BWX4491_xyl30m_1mM_25m | Hi-C |
| Figure S2A (row 1, column 6) | 315_Wang_HiC_BWX4491_xyl30m_1mM_30m | Hi-C |
| Figure S2A (row 2, column 1) | 304_Wang_HiC_BWX4493_xyl30m_1mM_00m | Hi-C |
| Figure S2A (row 2, column 2) | 305_Wang_HiC_BWX4493_xyl30m_1mM_10m | Hi-C |
| Figure S2A (row 2, column 3) | 306_Wang_HiC_BWX4493_xyl30m_1mM_15m | Hi-C |
| Figure S2A (row 2, column 4) | 307_Wang_HiC_BWX4493_xyl30m_1mM_20m | Hi-C |
| Figure S2A (row 2, column 5) | 308_Wang_HiC_BWX4493_xyl30m_1mM_25m | Hi-C |
| Figure S2A (row 2, column 6) | 309_Wang_HiC_BWX4493_xyl30m_1mM_30m | Hi-C |
| Figure S2A (row 3, column 1) | 316_Wang_HiC_BWX4492_xyl30m_1mM_00m | Hi-C |
| Figure S2A (row 3, column 2) | 317_Wang_HiC_BWX4492_xyl30m_1mM_10m | Hi-C |
| Figure S2A (row 3, column 3) | 318_Wang_HiC_BWX4492_xyl30m_1mM_15m | Hi-C |
| Figure S2A (row 3, column 4) | 319_Wang_HiC_BWX4492_xyl30m_1mM_20m | Hi-C |
| Figure S2A (row 3, column 5) | 320_Wang_HiC_BWX4492_xyl30m_1mM_25m | Hi-C |
| Figure S2A (row 3, column 6) | 321_Wang_HiC_BWX4492_xyl30m_1mM_30m | Hi-C |
| Figure S2B,C (left) | 301_Wang_HiC_BWX4476_1mM_060m | Hi-C |
| Figure S2B,C (middle) | 302_Wang_HiC_BWX4475_1mM_060m | Hi-C |
| Figure S2B,C (right) | 303_Wang_HiC_BWX4463_1mM_060m | Hi-C |
| Figure S3A | 303_Wang_HiC_BWX4463_1mM_060m | Hi-C |
| Figure S3B | 373_Wang_ChIPParB_BWX4462_1mM_060m | ChIP-seq |
| Figure S3B | 336_Wang_input_BWX4462_1mM_060m | WGS |
| Figure S5 | 303_Wang_HiC_BWX4463_1mM_060m | Hi-C |
| Figure S6 | 303_Wang_HiC_BWX4463_1mM_060m | Hi-C |
| Figure S7 | 303_Wang_HiC_BWX4463_1mM_060m | Hi-C |
| Figure S8 | 374_Wang_ChIPSMC_BWX3370 | ChIP-seq |
| Figure S8 | 375_Wang_input_BWX3370 | WGS |
| Figure S9A,B | 376_Wang_HiC_BWX4473_1mM_060m | Hi-C |
| Figure S10A,B | 302_Wang_HiC_BWX4475_1mM_060m | Hi-C |
| Figure S11A,B | 303_Wang_HiC_BWX4463_1mM_060m | Hi-C |
| Figure S12A (column 1) | 340_Wang_HiC_BWX4422 | Hi-C |
| Figure S12A (column 2) | 341_Wang_HiC_BWX4423 | Hi-C |
| Figure S12A (column 3) | 342_Wang_HiC_BWX4424 | Hi-C |
| Figure S12A (column 4) | 343_Wang_HiC_BWX4425 | Hi-C |
| Figure S12A (column 5) | 344_Wang_HiC_BWX4870 | Hi-C |
| Figure S12A (column 6) | 345_Wang_HiC_BWX4429 | Hi-C |
| Figure S12A (column 7) | 346_Wang_HiC_BWX4885 | Hi-C |
| Figure S12B (column 1) | 347_Wang_HiC_BWX4891 | Hi-C |
| Figure S12B (column 2) | 348_Wang_HiC_BWX5066_OIPTG | Hi-C |
| Figure S13A (top column 1) | 302_Wang_HiC_BWX4475_1mM_060m | Hi-C |
| Figure S13A (top column 2) | 349_Wang_HiC_BWX4475_1mM_090m | Hi-C |
| Figure S13A (top column 3) | 350_Wang_HiC_BWX4475_1mM_120m | Hi-C |
| Figure S13A (bottom column 1) | 351_Wang_HiC_BWX4515_1mM_060m | Hi-C |
| Figure S13A (bottom column 2) | 352_Wang_HiC_BWX4515_1mM_090m | Hi-C |
| Figure S13A (bottom column 3) | 353_Wang_HiC_BWX4515_1mM_120m | Hi-C |
| Figure S13B (top column 1) | 326_Wang_HiC_BWX4883_1mM_060m | Hi-C |
| Figure S13B (top column 2) | 354_Wang_HiC_BWX4883_1mM_090m | Hi-C |
| Figure S13B (top column 3) | 355_Wang_HiC_BWX4883_1mM_120m | Hi-C |
| Figure S13B (bottom column 1) | 328_Wang_HiC_BWX4892_1mM_060m | Hi-C |
| Figure S13B (bottom column 2) | 356_Wang_HiC_BWX4892_1mM_090m | Hi-C |
| Figure S13B (bottom column 3) | 357_Wang_HiC_BWX4892_1mM_120m | Hi-C |
| Figure S14A (column 1) | 366_Wang_input_BWX4462_OIPTG | WGS |
| Figure S14A (column 2) | 336_Wang_input_BWX4462_1mM_060m | WGS |
| Figure S14A (column 3) | 337_Wang_input_BWX4462_1mM_090m | WGS |
| Figure S14A (column 4) | 338_Wang_input_BWX4462_1mM_120m | WGS |
| Figure S15A (left) | 322_Wang_HiC_BWX4462_1mM_060m | Hi-C |
| Figure S15A (middle) | 331_Wang_HiC_BWX4462_1mM_090m | Hi-C |
| Figure S15A (right) | 332_Wang_HiC_BWX4462_1mM_120m | Hi-C |
| Figure S15B (left) | 358_Wang_HiC_BWX5132_1mM_060m | Hi-C |
| Figure S15B (middle) | 359_Wang_HiC_BWX5132_1mM_090m | Hi-C |
| Figure S15B (right) | 360_Wang_HiC_BWX5132_1mM_120m | Hi-C |

| **figure** | **sample** | **data type** |
| --- | --- | --- |
| Figure S15C (top) | 322_Wang_HiC_BWX4462_1mM_060m | Hi-C |
| Figure S15C (center) | 358_Wang_HiC_BWX5132_1mM_060m | Hi-C |
| Figure S15C (bottom) | 331_Wang_HiC_BWX4462_1mM_090m | Hi-C |
| Figure S16A (left) | 330_Wang_HiC_BWX5066_1mM_060m | Hi-C |
| Figure S16A (middle) | 361_Wang_HiC_BWX5066_1mM_090m | Hi-C |
| Figure S16A (right) | 362_Wang_HiC_BWX5066_1mM_120m | Hi-C |
| Figure S16B (left) | 367_Wang_ChIPSMC_BWX5066rep2_1mM_060m | ChIP-seq |
| Figure S16B (middle) | 368_Wang_ChIPSMC_BWX5066rep2_1mM_090m | ChIP-seq |
| Figure S16B (right) | 369_Wang_ChIPSMC_BWX5066rep2_1mM_120m | ChIP-seq |
| Figure S16B (left) | 370_Wang_input_BWX5066rep2_1mM_060m | WGS |
| Figure S16B (middle) | 371_Wang_input_BWX5066rep2_1mM_090m | WGS |
| Figure S16B (right) | 372_Wang_input_BWX5066rep2_1mM_120m | WGS |
| Figure S17A (left) | 363_Wang_HiC_BWX4359_1mM_060m | Hi-C |
| Figure S17A (middle) | 364_Wang_HiC_BWX4359_1mM_090m | Hi-C |
| Figure S17A (right) | 365_Wang_HiC_BWX4359_1mM_120m | Hi-C |
| Figure S18B | 01_Rudnerlab_HindIII_HiC_PY79<sup>4</sup> | Hi-C |

**Supplemental Table S2.** Bacterial strains used in this study.

| strain | genotype | reference | figure |
| --- | --- | --- | --- |
| BWX3221 | *parS*Δ9 (*loxP-spec-loxP*), -94° *parS* (*loxP-kan-loxP*) | 4 | S1B |
| BWX3370 | *parS*Δ9 no a.b., -1° *parS* | 8 | S8 |
| BWX3377 | *parS*Δ9 no a.b., -59° *parS* (*loxP-kan-loxP*) | 8 | S1B |
| BWX4359 | *yhdG::Phyperspank-(optRBS)-sirA* (phleo) | this study | S17A |
| BWX4422 | *parS*Δ9 no a.b., -27° *parS*, -59° *parS* (*loxP-kan-loxP*) | this study | S12A |
| BWX4423 | *parS*Δ9 no a.b., -27° *parS*, -94° *parS* (*loxP-kan-loxP*) | this study | S12A |
| BWX4424 | *parS*Δ9 no a.b., -27° *parS*, -117° *parS* (*loxP-kan-loxP*) | this study | S12A |
| BWX4425 | *parS*Δ9 no a.b., -27° *parS*, -153° *parS* (*loxP-kan-loxP*) | this study | S12A |
| BWX4428 | *parS*Δ9 no a.b., -59° *parS* no a.b., -94° *parS* (*loxP-kan-loxP*) | this study | S1B |
| BWX4429 | *parS*Δ9 no a.b., -59° *parS* no a.b., -117° *parS* (*loxP-kan-loxP*) | this study | S12A |
| BWX4462 | *parS*Δ9 no a.b., -27° *parS*, -59° *parS* (*loxP-kan-loxP*), *yhdG::Phyperspank-(optRBS)-sirA* (phleo) | this study | 3AC, 4ABC, S3B, S14AC, S15AC |
| BWX4463 | *parS*Δ9 no a.b., -59° *parS* no a.b., -94° *parS* (*loxP-kan-loxP*), *yhdG::Phyperspank-(optRBS)-sirA* (phleo) | this study | 1C, 2AC, S2BC, S3A, S5, S6, S7, S11AB |
| BWX4473 | *parS*Δ9 no a.b., *yhdG::Phyperspank-(optRBS)-sirA* (phleo) | this study | S9AB |
| BWX4475 | *parS*Δ9 no a.b., -59° *parS* (*loxP-kan-loxP*), *yhdG::Phyperspank-(optRBS)-sirA* (phleo) | this study | 1C, S2BC, S10AB, S13A |
| BWX4476 | *parS*Δ9 no a.b., -94° *parS* no a.b., *yhdG::Phyperspank-(optRBS)-sirA* (phleo) | this study | 1C, S2BC |
| BWX4479 | *parS*Δ9 no a.b., -27° *parS*, -94° *parS* (*loxP-kan-loxP*), *yhdG::Phyperspank-(optRBS)-sirA* (phleo) | this study | 3A |
| BWX4480 | *parS*Δ9 no a.b., -27° *parS*, -117° *parS* (*loxP-kan-loxP*), *yhdG::Phyperspank-(optRBS)-sirA* (phleo) | this study | 3A |
| BWX4481 | *parS*Δ9 no a.b., -27° *parS*, -153° *parS* (*loxP-kan-loxP*), *yhdG::Phyperspank-(optRBS)-sirA* (phleo) | this study | 3A |
| BWX4482 | *parS*Δ9 no a.b., -59° *parS* no a.b., -117° *parS* (*loxP-kan-loxP*), *yhdG::Phyperspank-(optRBS)-sirA* (phleo) | this study | 3A |
| BWX4491 | *parS*Δ9 no a.b., -59° *parS* no a.b., Δ*parB*(Δ*parS*) (*loxP-spec-loxP*), *yvbJ::Pspank-(optRBS)-parB*(Δ*parS*) (cat), *yhdG::Pxyl-(optRBS)-sirA* (phleo) | this study | 1D, S2A |
| BWX4492 | *parS*Δ9 no a.b., -94° *parS* no a.b., Δ*parB*(Δ*parS*) (*loxP-spec-loxP*), *yvbJ::Pspank-(optRBS)-parB*(Δ*parS*) (cat), *yhdG::Pxyl-(optRBS)-sirA* (phleo) | this study | 1D, S2A |
| BWX4493 | *parS*Δ9 no a.b., -59° *parS* no a.b., -94° *parS* (*loxP-kan-loxP*), Δ*parB*(Δ*parS*) (*loxP-spec-loxP*), *yvbJ::Pspank-(optRBS)-parB*(Δ*parS*) (cat), *yhdG::Pxyl-(optRBS)-sirA* (phleo) | this study | 1D, S2A |
| BWX4515 | *parS*Δ9 no a.b., -91° *parS* (*loxP-kan-loxP*), *yhdG::Phyperspank-(optRBS)-sirA* (phleo) | this study | S13A |
| BWX4547 | *yycR* (-7°)::*tetO48* (cat), *ycgO::PftsW-tetR-cfp* (phleo), *yhdG::Phyperspank-(optRBS)-sirA* (erm) | this study | 4C |
| BWX4870 | *parS*Δ9 no a.b., -59° *parS* no a.b., -91° *parS* (*loxP-kan-loxP*) | this study | S12A |
| BWX4883 | *parS*Δ9 no a.b., -59° *parS* no a.b., -91° *parS* (*loxP-kan-loxP*), *yhdG::Phyperspank-(optRBS)-sirA* (phleo) | this study | 3A, S13B |
| BWX4885 | *parS*Δ9 no a.b., -91° *parS* no a.b., -117° *parS* (*loxP-kan-loxP*) | this study | S12A |
| BWX4891 | *parS*Δ9 no a.b., -59° *parS* no a.b., -91° *parS* no a.b., -117° *parS* (*loxP-kan-loxP*) | this study | S12B |
| BWX4892 | *parS*Δ9 no a.b., -91° *parS* no a.b., -117° *parS* (*loxP-kan-loxP*), *yhdG::Phyperspank-(optRBS)-sirA* (phleo) | this study | 3A, S13B |
| BWX4927 | *parS*Δ9 no a.b., -59° *parS* no a.b., -91° *parS* no a.b., -117° *parS* (*loxP-kan-loxP*), *yhdG::Phyperspank-(optRBS)-sirA* (phleo) | this study | 3B |
| BWX5066 | *parS*Δ9 no a.b., -27° *parS*, -59° *parS* no a.b., -91° *parS* (*loxP-kan-loxP*), *yhdG::Phyperspank-(optRBS)-sirA* (phleo) | this study | 3B, S12B, S16AB |
| BWX5132 | *parS*Δ9 no a.b., -27° *parS*, -59° *parS* (*loxP-kan-loxP*), *amyE::Phyperspank-(optRBS)-smc* (spec), *yhdG::Phyperspank-(optRBS)-scpAB* (phleo), *yvbJ::Phyperspank-(optRBS)-sirA* (erm) | this study | S15BC |
| PY79 | wild-type | 1 | S18B |
| BNS1615 | *parS*Δ7: *spo0J* (*parS*Δ), *yycG* (*parS*Δ), *rocR* (*parS*Δ), *cotF* (*parS*Δ), *metS* (*parS*Δ), *ybbC*(*parS*Δ), *ydaD*(*parS*Δ) | 4 | |
| BNS1657 | *parS*Δ8: *parB* (*parS*Δ), *yycG* (*parS*Δ), *rocR* (*parS*Δ), *cotF* (*parS*Δ), *metS* (*parS*Δ), *ybbC*(*parS*Δ), *ydaD*(*parS*Δ), *nfrA*(*parS*Δ) | 17 | |
| BWX811 | *yycR* (-7°)::*tetO48* (cat), *ycgO::PftsW-tetR-cfp* (phleo) | 18 |  |
| BWX2761 | *parS*Δ8, Δ*parB* (Δ*parS*)::spec | 4 | |
| BWX3198 | *parS*Δ8, +91° *yhaX* (Δ*parS*) (*loxP*-*kan*-*loxP*) | 4 | |
| BWX3212 | *parS*Δ9 no a.b. | 4 | |
| BWX3268 | *parS*Δ9 no a.b., -27° *parS* | 8 | |
| BWX3270 | *parS*Δ9 no a.b., -94° no a.b. | 8 | |
| BWX3381 | *parS*Δ9 no a.b., -117° *parS* (*loxP*-*kan*-*loxP*) | 8 | |
| BWX3383 | *parS*Δ9 no a.b., -153° *parS* (*loxP*-*kan*-*loxP*) | 8 | |

**Supplemental Table S3.** Plasmids used in this study.

| plasmid | description | reference |
| --- | --- | --- |
| pJW005 | *yhdG::Phyperspank-(optRBS)-sirA* (*phleo*) | 3 |
| pWX512 | *amyE::Phyperspank-(optRBS)-smc* (*spec*) | this study |
| pWX722 | *yvbJ::Pspank-(optRBS)-parB*(Δ*parS*) (*cat*) | 19 |
| pWX777 | *yhdG::Pxyl-(optRBS)-sirA* (*phleo*) | this study |
| pWX778 | *yhdG::Phyperspank-(optRBS)-scpAB* (*phleo*) | this study |
| pWX788 | *yhdG::Phyperspank-(optRBS)-sirA* (*erm*) | this study |

**Supplemental Table S4.**
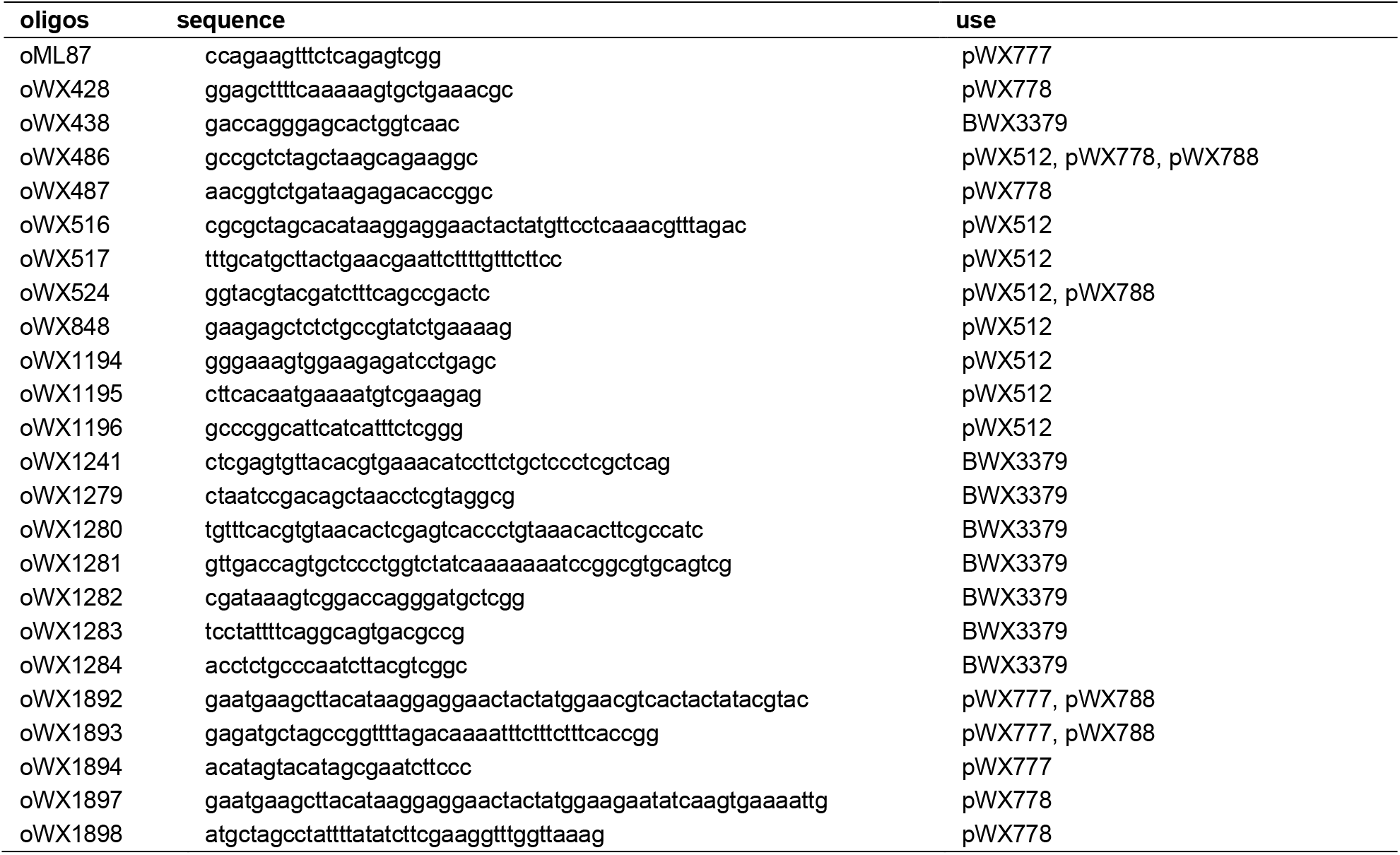
Oligonucleotides used in this study.

## References

1. Hirano, T. Condensins: Universal organizers of chromosomes with diverse functions. Genes Dev. 26, 1659–1678 (2012).

2. Yatskevich, S., Rhodes, J. & Nasmyth, K. Organization of Chromosomal DNA by SMC Complexes. Annu. Rev. Genet. 53, 445–482 (2019).

3. Wang, X., Brandão, H. B., Le, T. B. K., Laub, M. T. & Rudner, D. Z. Bacillus subtilis SMC complexes juxtapose chromosomal arms as they travel from origin to terminus. Science. 355, 524–527 (2017).

4. Tran, N. T., Laub, M. T. & Le, T. B. K. SMC Progressively Aligns Chromosomal Arms in Caulobacter crescentus but Is Antagonized by Convergent Transcription. Cell Rep. 20, 2057–2071 (2017).

5. Terakawa, T. et al. The condensin complex is a mechanochemical motor that translocates along DNA. Science. 358, 672–676 (2017).

6. Wang, X. et al. In vivo evidence for ATPase-dependent DNA translocation by the Bacillus subtilis SMC condensin complex. Mol. Cell 71, 841–847 (2018).

7. Ganji, M. et al. Real-time imaging of DNA loop extrusion by condensin. Science. 360, 102–105 (2018).

8. Kim, Y., Shi, Z., Zhang, H., Finkelstein, I. J. & Yu, H. Human cohesin compacts DNA by loop extrusion. Science. 366, 1345–1349 (2019).

9. Davidson, I. F. et al. Human cohesin compacts DNA by loop extrusion. Science. 366, 1338–1345 (2019).

10. Kong, M. et al. Human condensin I and II drive extensive ATP–dependent compaction of nucleosome–bound DNA. Mol. Cell 79, 1–16 (2020).

11. Golfier, S., Quail, T., Kimura, H. & Brugués, J. Cohesin and condensin extrude DNA loops in a cell-cycle dependent manner. Elife 9, e53885 (2020).

12. Kim, E., Kerssemakers, J., Shaltiel, I. A., Haering, C. H. & Dekker, C. DNA-loop extruding condensin complexes can traverse one another. Nature 579, 438–442 (2020).

13. Sullivan, N. L., Marquis, K. A. & Rudner, D. Z. Recruitment of SMC by ParB-parS organizes the origin region and promotes efficient chromosome segregation. Cell 137, 697–707 (2009).

14. Gruber, S. & Errington, J. Recruitment of condensin to replication origin regions by ParB/SpoOJ promotes chromosome segregation in B. subtilis. Cell 137, 685–696 (2009).

15. Minnen, A. et al. Control of Smc coiled coil architecture by the ATPase heads facilitates targeting to chromosomal ParB/parS and release onto flanking DNA. Cell Rep. 14, 2003–2016 (2016).

16. Böhm, K. et al. Chromosome organization by a conserved condensin-ParB system in the actinobacterium Corynebacterium glutamicum. Nat. Commun. 11, 1–17 (2020).

17. Lioy, V., Junier, I., Lagage, V., Vallet, I. & Boccard, F. Distinct activities of bacterial condensins for chromosome management in Pseudomonas aeruginosa. BioRxiv 1–41 (2020). doi:10.1101/2020.05.18.101659

18. Fudenberg, G., Abdennur, N., Imakaev, M., Goloborodko, A. & Mirny, L. A. Emerging evidence of chromosome folding by loop extrusion. Cold Spring Harb. Symp. Quant. Biol. 82, 45–55 (2017).

19. Banigan, E. J. & Mirny, L. A. Loop extrusion: theory meets single-molecule experiments. Curr. Opin. Cell Biol. 64, 124–138 (2020).

20. Livny, J., Yamaichi, Y. & Waldor, M. K. Distribution of centromere-like parS sites in bacteria: Insights from comparative genomics. J. Bacteriol. 189, 8693–8703 (2007).

21. Lioy, V. S. et al. Multiscale Structuring of the E. coli Chromosome by Nucleoid-Associated and Condensin Proteins. Cell 172, 771–783 (2018).

22. Mäkelä, J. & Sherratt, D. J. Organization of the Escherichia coli Chromosome by a MukBEF Axial Core. Mol. Cell 78, 250–260 (2020).

23. Alipour, E. & Marko, J. F. Self-organization of domain structures by DNA-loop-extruding enzymes. Nucleic Acids Res. 40, 11202–11212 (2012).

24. Breier, A. M. & Grossman, A. D. Whole-genome analysis of the chromosome partitioning and sporulation protein Spo0J (ParB) reveals spreading and origin-distal sites on the Bacillus subtilis chromosome. Mol. Microbiol. 64, 703–718 (2007).

25. Wang, X. et al. Condensin promotes the juxtaposition of DNA flanking its loading site in Bacillus subtilis. Genes Dev. 29, 1661–1675 (2015).

26. Marbouty, M. et al. Condensin- and replication-mediated bacterial chromosome folding and origin condensation revealed by Hi-C and super-resolution imaging. Mol. Cell 59, 588–602 (2015).

27. Wagner, J. K., Marquis, K. A. & Rudner, D. Z. SirA enforces diploidy by inhibiting the replication initiator DnaA durina snore formation in Bacillus subtilis. Mol. Microbiol. 73, 963–974 (2009).

28. Le, T. B. K., Imakaev, M. V., Mirny, L. A. & Laub, M. T. High-resolution mapping of the spatial organization of a bacterial chromosome. Science. 342, 731–734 (2013).

29. Brandão, H. B. et al. RNA polymerases as moving barriers to condensin loop extrusion. Proc. Natl. Acad. Sci. U. S. A. 116, 20489–20499 (2019).

30. Banigan, E. J., van den Berg, A. A., Brandão, H. B., Marko, J. F. & Mirny, L. A. Chromosome organization by one-sided and two-sided loop extrusion. Elife 9, e53558 (2020).

31. Miermans, C. A. & Broedersz, C. P. Bacterial chromosome organization by collective dynamics of SMC condensins. J. R. Soc. Interface 15, 20180495 (2018).

32. Wilhelm, L. et al. SMC condensin entraps chromosomal DNA by an ATP hydrolysis dependent loading mechanism in Bacillus subtilis. Elife 4, e06659 (2015).

33. Sueoka, N. & Yoshikawa, H. The chromosome of Bacillus subtilis. I. Theory of marker frequency analysis. Genetics 52, 747–757 (1965).

34. Vazquez Nunez, R., Ruiz Avila, L. B. & Gruber, S. Transient DNA Occupancy of the SMC Interarm Space in Prokaryotic Condensin. Mol. Cell 75, 209–223 (2019).

35. He, Y. et al. Statistical mechanics of chromosomes: in vivo and in silico approaches reveal high-level organization and structure arise exclusively through mechanical feedback between loop extruders and chromatin substrate properties. Nucleic Acids Res. 1, gkaa871 (2020).

36. Liu, Y. et al. Very Fast CRISPR on Demand. Science. 368, 1265–1269 (2020).

37. Arnould, C. et al. Loop extrusion as a mechanism for DNA Double-Strand Breaks repair foci formation. (2020). doi:10.1101/2020.02.12.945311

38. Li, K., Bronk, G., Kondev, J. & Haber, J. E. Yeast ATM and ATR kinases use different mechanisms to spread histone H2A phosphorylation around a DNA double-strand break. Proc. Natl. Acad. Sci. U. S. A. 117, 21354–21363 (2020).

39. Wang, C. Y., Colognori, D., Sunwoo, H., Wang, D. & Lee, J. T. PRC1 collaborates with SMCHD1 to fold the X-chromosome and spread Xist RNA between chromosome compartments. Nat. Commun. 10, 1–18 (2019).

40. Anderson, E. C. et al. X Chromosome Domain Architecture Regulates Caenorhabditis elegans Lifespan but Not Dosage Compensation. Dev. Cell 51, 192–207 (2019).

41. Heger, P., Marin, B., Bartkuhn, M., Schierenberg, E. & Wiehe, T. The chromatin insulator CTCF and the emergence of metazoan diversity. Proc. Natl. Acad. Sci. U. S. A. 109, 17507–17512 (2012).

## Supplemental References

1 Youngman, P. J., Perkins, J. B. & Losick, R. Genetic transposition and insertional mutagenesis in Bacillus subtilis with Streptococcus faecalis transposon Tn917. Proc Natl Acad Sci U S A 80, 2305–2309 (1983).

2 Harwood, C. R. & Cutting, S. M. Molecular Biological Methods for Bacillus. (Wiley, 1990).

3 Wagner, J. K., Marquis, K. A. & Rudner, D. Z. SirA enforces diploidy by inhibiting the replication initiator DnaA during spore formation in Bacillus subtilis. Mol Microbiol 73, 963–974, doi:MMI6825 [pii] 10.1111/j.1365-2958.2009.06825.x (2009).

4 Wang, X. et al. Condensin promotes the juxtaposition of DNA flanking its loading site in Bacillus subtilis. Genes Dev 29, 1661–1675, doi:10.1101/gad.265876.115 (2015).

5 Lindow, J. C., Kuwano, M., Moriya, S. & Grossman, A. D. Subcellular localization of the Bacillus subtilis structural maintenance of chromosomes (SMC) protein. Mol Microbiol 46, 997–1009 (2002).

6 Lin, D. C., Levin, P. A. & Grossman, A. D. Bipolar localization of a chromosome partition protein in Bacillus subtilis. Proc Natl Acad Sci U S A 94, 4721–4726 (1997).

7 Fujita, M. Temporal and selective association of multiple sigma factors with RNA polymerase during sporulation in Bacillus subtilis. Genes Cells 5, 79–88, doi:gtc307 [pii] (2000).

8 Wang, X., Brandao, H. B., Le, T. B., Laub, M. T. & Rudner, D. Z. Bacillus subtilis SMC complexes juxtapose chromosome arms as they travel from origin to terminus. Science 355, 524–527, doi:10.1126/science.aai8982 (2017).

9 Meeske, A. J. et al. MurJ and a novel lipid II flippase are required for cell wall biogenesis in Bacillus subtilis. Proc Natl Acad Sci U S A 112, 6437–6442, doi:10.1073/pnas.1504967112 (2015).

10 Harris, C. R. et al. Array programming with NumPy. Nature 585, 357–362, doi:10.1038/s41586-020-2649-2 (2020).

11 Banigan, E. J., van den Berg, A. A., Brandao, H. B., Marko, J. F. & Mirny, L. A. Chromosome organization by one-sided and two-sided loop extrusion. eLife 9, doi:10.7554/eLife.53558 (2020).

12 Brandao, H. B. et al. RNA polymerases as moving barriers to condensin loop extrusion. Proc Natl Acad Sci U S A 116, 20489–20499, doi:10.1073/pnas.1907009116 (2019).

13 Imakaev, M., Goloborodko, A. & Brandao, H. B. in Zenodo (2019). http://doi.org/10.5281/zenodo.3579473

14 Eastman, P. et al. OpenMM 7: Rapid development of high performance algorithms for molecular dynamics. PLoS Comput Biol 13, e1005659, doi:10.1371/journal.pcbi.1005659 (2017).

15 Virtanen, P. et al. SciPy 1.0: fundamental algorithms for scientific computing in Python. Nat Methods 17, 261–272, doi:10.1038/s41592-019-0686-2 (2020).

16 Kim, E., Kerssemakers, J., Shaltiel, I. A., Haering, C. H. & Dekker, C. DNA-loop extruding condensin complexes can traverse one another. Nature 579, 438–442, doi:10.1038/s41586-020-2067-5 (2020).

17 Sullivan, N. L., Marquis, K. A. & Rudner, D. Z. Recruitment of SMC by ParB-parS organizes the origin region and promotes efficient chromosome segregation. Cell 137, 697–707, doi:10.1016/j.cell.2009.04.044 (2009).

18 Wang, X., Montero Llopis, P. & Rudner, D. Z. Bacillus subtilis chromosome organization oscillates between two distinct patterns. Proc Natl Acad Sci U S A, doi:10.1073/pnas.1407461111 (2014).

19 Wang, X. et al. In Vivo Evidence for ATPase-Dependent DNA Translocation by the Bacillus subtilis SMC Condensin Complex. Mol Cell 71, 841–847 e845, doi:10.1016/j.molcel.2018.07.006 (2018).

